# Heme-dependent siderophore utilization promotes iron-restricted growth of the *Staphylococcus aureus hemB* small colony variant

**DOI:** 10.1101/2021.06.29.450315

**Authors:** Izabela Z. Batko, Ronald S. Flannagan, Veronica Guariglia-Oropeza, Jessica R. Sheldon, David E. Heinrichs

## Abstract

The ability to acquire iron is essential for *Staphylococcus aureus* to cause infection. Respiration deficient *S. aureus* small colony variants (SCVs) frequently cause persistent infections, which necessitates they too acquire iron. How SCVs obtain iron remains unknown and so here we addressed this outstanding question by creating a stable *hemB* mutant in *S. aureus* USA300 strain LAC. The mutant, auxotrophic for hemin, was assessed for its ability to grow under iron-restriction and with various iron sources. The *hemB* SCV utilizes exogenously supplied heme but was attenuated for growth under conditions of iron starvation. RNA-seq analyses showed that both WT *S. aureus* and the *hemB* mutant sense and respond to iron starvation, however, growth assays show that the *hemB* mutant is defective for siderophore-mediated iron acquisition. Indeed, the *hemB* SCV demonstrates limited utilization of endogenous staphyloferrin B or exogenously provided staphyloferrin A, Desferal, and epinephrine, which enabled the SCV to sustain only minimal growth in iron deplete media. Direct measurement of intracellular ATP in *hemB* and WT *S. aureus* revealed that both strains can generate comparable levels of ATP during exponential growth suggesting defects in ATP production cannot account for the inability to efficiently utilize siderophores. Defective siderophore utilization by *hemB* bacteria was also evident *in vivo*. Indeed, the administration of Desferal failed to promote *hemB* bacterial growth *in vivo*, in contrast to WT, in every organ analyzed except for the murine kidney where growth was enhanced. In support of the hypothesis that *S. aureus* accesses heme in kidney abscesses, *in vitro* analyses revealed that increased heme availability enables *hemB* bacteria to utilize siderophores for growth when iron availability is restricted. Taken together, our data support the conclusion that heme is not only used as an iron source itself, but as a nutrient that promotes utilization of siderophore-iron complexes.

**Author summary:** *S. aureus* small colony variants (SCVs) represent a difficult to treat subpopulation of bacteria that are associated with chronic recurrent infection and worsened clinical outcome. Indeed, SCVs persist within the host despite appropriate administration of antibiotics. This study yields insight into the mechanisms by which *S. aureus* SCVs acquire iron which, during infection of a host, is a difficult-to-acquire metal nutrient. Under heme limited conditions, *hemB S. aureus* is severely impaired for siderophore-dependent growth and, in agreement, murine infection indicates that hemin-deficient SCVs meet their nutritional requirement for iron through utilization of heme *in vivo*. Importantly, we demonstrate that *hemB* SCVs rely upon heme as a nutrient to promote siderophore utilization. Therefore, perturbation of heme biosynthesis and/or utilization represents a viable to strategy to mitigate the ability of SCV bacteria to acquire siderophore-bound iron during infection.

## Introduction

The notorious pathogen *Staphylococcus aureus* is a Gram-positive bacterium that can colonize virtually every tissue of the human body. *S. aureus* infections range from mild skin and soft tissue infections to life-threatening conditions such as pneumonia, endocarditis, and osteomyelitis [1, 2]. Importantly, *S. aureus* is a significant cause of morbidity and mortality, which is emphasized by reports that invasive *S. aureus* infections contribute to more deaths in the United States than HIV [3–6]. The success of *S. aureus* as a pathogen is dependent on its ability to sense the host environment, evade the host immune system, alter its metabolism appropriately, and acquire or synthesize essential nutrients [7–16].

Iron is an essential nutrient for most lifeforms, including *S. aureus* [17]. However, hosts strictly regulate iron metabolism and actively sequester iron to limit its availability, thereby mitigating the potentially harmful effects of free iron and helping to curtail microbial growth [18–20]. The process of actively starving pathogens of essential nutrients, such as iron, to promote bacteriostasis is termed “nutritional immunity” [21]. Successful pathogens such as *S. aureus* have evolved highly specialized iron acquisition mechanisms that enable bacteria to circumvent host iron restriction mechanisms [22].

The ability of *S. aureus* to acquire iron within iron restricted environments requires coordinated expression of iron acquisition genes that are regulated by the ferric uptake regulator (Fur) [23]. The iron acquisition strategies employed by *S. aureus* have been studied at length and recently reviewed [24, 25]. The iron-regulated surface determinant (Isd) pathway [26–30] is a high-affinity heme iron acquisition system expressed by *S. aureus*, the importance of which is evident at even very low concentrations (<50 nM) of heme [31]. *S. aureus* can also utilize two endogenously synthesized siderophores, staphyloferrin A (SA) and staphyloferrin B (SB), as well as so-called xenosiderophores, that originate from other bacteria, to capture trace amounts of non-heme iron; reviewed in [24,25,32].

*S. aureus* small colony variants (SCVs) represent a slow-growing subpopulation of *S. aureus* that display increased resistance to antibiotics and persistence within the host [33–36]. The clinical relevance of *S. aureus* SCVs is underscored by the routine isolation of SCVs from patients with chronic *S. aureus* infections [33–36]. *S. aureus* SCVs are highly prevalent among cystic fibrosis patients who often experience chronic recurrent lung infections that are associated with worse clinical outcomes [37–39]. Clinically isolated *S. aureus* SCVs often demonstrate auxotrophy for hemin (ferric chloride heme) and display defects in bacterial respiration [33, 34]. Heme is a co-factor found in the terminal oxidases encoded by the genes *cydAB* and *qoxABCD* in *S. aureus* and inactivation of these genes results in SCV-like growth [40]. The importance of these oxidases to *S. aureus* fitness is underscored by the finding that oxidase inactivation causes significant defects in organ colonization in a murine model of systemic infection [40]. Several phenotypes have been ascribed to SCV bacteria including altered central metabolism, reduced toxin expression, increased resistance to antimicrobial peptides and antibiotics, persistence within host cells and, notably, a reduced growth rate [36]. Remarkably, how respiration defects affect iron acquisition in *S. aureus* has not been explored. Since SCVs arise and persist during infection it stands to reason that these bacteria acquire iron within the host.

Here we explored how a stable *S. aureus hemB* SCV grows under iron restricted conditions *in vitro* and *in vivo*. Importantly, we demonstrate that under conditions of iron limitation, versus iron replete conditions, SCV growth is severely constrained, and this is due to impaired iron-siderophore utilization, a defect also evident during infection of the host using a systemic model of infection in mice. We further provide evidence that the inability of *hemB* mutant bacteria to utilize siderophores is dependent on the availability of heme and that utilization of exogenous heme permits siderophore-dependent growth of iron-starved *hemB* SCVs. Our results provide valuable insights into the iron acquisition strategies employed by *S. aureus* SCVs and indicate that perturbation of heme utilization could render SCVs unable to grow under iron limited conditions such as those presented by the host.

## Results

### Analysis of iron-restricted growth of a *S. aureus* USA300 *hemB* mutant

At the outset of this study, we generated a stable SCV in the MRSA strain USA300-LAC (hereafter referred to as USA300) by mutating the *hemB* gene; naturally occurring SCVs often contain mutations in *hemB*, which obviate heme biosynthesis leading to a respiration defective phenotype [41, 42]. The USA300 *hemB* mutant demonstrated characteristics expected of an SCV, including a small colony morphology on tryptic soy agar and decreased toxin production, the latter typified by low hemolytic activity on blood agar (Fig S1). The USA300 *hemB* mutant phenotype was fully complemented if the *hemB* gene was provided *in trans* and, moreover, supplementation of the growth medium with the metabolite hemin also complemented the growth phenotype, although not to WT levels (Fig S1).

To begin to investigate the iron starvation response of the *hemB* mutant, we demonstrated that while the mutant grew poorly in TSB as compared to control strains (Fig 1A), it did not grow at all in the minimal medium TMS (Fig 1B and 1C), unless hemin was added (Fig 1C). These data confirm the growth requirement of the *hemB* mutant for heme [42], and also suggest that TSB contains traces of heme that allows minimal growth in that medium. Use of TMS, which is devoid of hemin, allowed us to control the amount of hemin that was available to the bacteria, as it could be provided exogenously. Of note, hemin supplementation up to approximately 2 µM increased bacterial growth (Fig S1) but above these concentrations we observed that it inhibited growth, as has been demonstrated previously [43–45]. Using this information, we established a condition that could support *hemB* mutant growth without fully complementing the SCV phenotype. These analyses revealed that as little as 0.4 µM hemin enabled *hemB S. aureus* to grow but not to WT levels (Fig 1C). Next, human transferrin (hTf) was used as an additive to the growth medium to restrict iron availability; this was aptly demonstrated by its ability to completely restrict the growth of a staphyloferrin-deficient (note that staphyloferrins [siderophores] can remove iron from hTf) strain of *S. aureus* at a minimum concentration of 1.5 µM hTf (Fig S2A). We observed that this concentration of hTf reduced *hemB S. aureus* growth as compared to a no hTf condition and the reduced growth could be corrected by supplementation of the medium with ferrous ammonium sulfate (Fe) to saturate the hTf (Fig 1D); Fe supplementation also allowed growth of the staphyloferrin-deficient strain in the presence of hTf (Fig S2A). Under the same culture conditions, WT *S. aureus* and the complemented *hemB* mutant grew robustly (Fig S2B and S2C). Together, these observations led us to conclude that iron restriction could significantly alter growth of the *S. aureus hemB* SCV, in comparison to WT, and that the *hemB* mutant must be impaired for iron acquisition.

**Figure 1.**
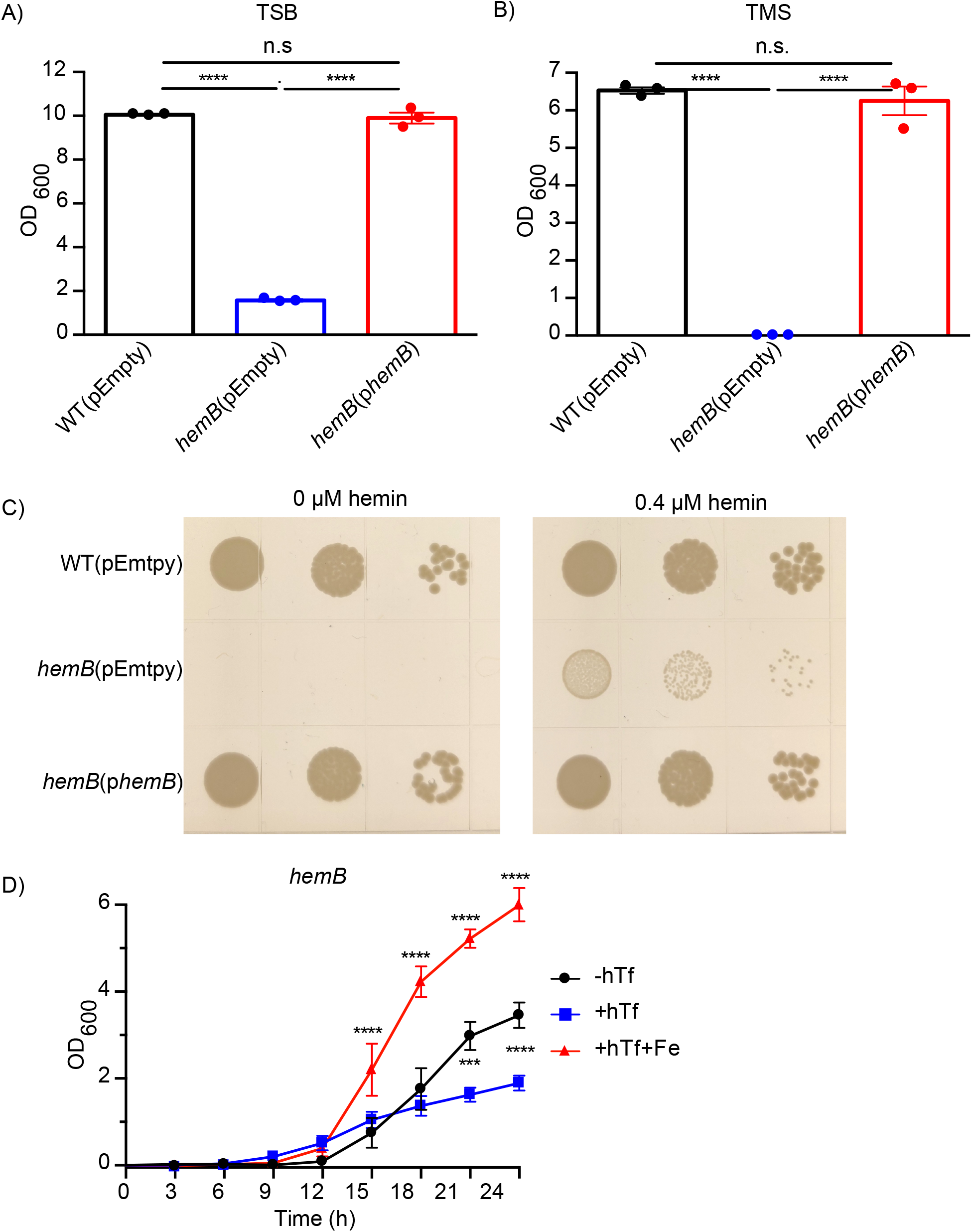
Growth of *S. aureus hemB* SCV in rich and minimal media. Growth of *S. aureus* USA300 WT(pEmpty), *hemB*(pEmpty), and *hemB*(p*hemB*) cultured in **(A)** tryptic soy broth (TSB) or **(B)** Tris minimal succinate (TMS) for 24 h. Data are plotted as mean ± SEM, three biological replicates. **** *p* ≤ 0.0001, one-way ANOVA with Tukey’s multiple comparisons. **(C)** Growth of *S. aureus* USA300 WT(pEmpty), *hemB*(pEmpty), and *hemB*(p*hemB*) on TMS agar supplemented with either 0 µM (left panel) or 0.4 µM (right panel) hemin after 48 h incubation at 37°C. **(D)** Growth over 24 h at 37°C of the *hemB* mutant cultured in TMS supplemented with 0.4 µM hemin and either no human transferrin (-hTf), 1.5 µM human transferrin (+hTf), or 1.5 µM human transferrin and 10 µM ferric ammonium sulfate (+hTf+Fe). Data are plotted as mean ± SEM, five biological replicates. \*\*\**p* ≤ 0.001, **** *p* ≤ 0.0001, two-way ANOVA with Dunnett’s multiple comparisons.

### The *S. aureus hemB* mutant responds to iron starvation

The transcriptional profile of *S. aureus* SCVs varies greatly from WT *S. aureus* [46–49], but it is unknown whether SCVs differentially express genes involved in Fur-regulated iron acquisition systems. Therefore, RNA-seq was performed to profile changes in gene expression between iron-starved WT *S. aureus* and the iron-starved *hemB* SCV. To that end, wild-type and *hemB* bacteria were cultured under iron-restricted growth conditions until each strain had achieved mid log phase. At this time the cultures were left untreated or spiked with iron and RNA was extracted after 1h. This analysis enabled assessment of the ability of *hemB S. aureus* to rapidly respond to fluctuations in extracellular iron. As expected of *S. aureus* SCVs, our analysis revealed that gene expression of the *hemB* mutant grown in TMS supplemented with 0.4 µM hemin vastly differs from that of WT, with 310 genes upregulated and 462 genes downregulated in the SCV, compared to WT, irrespective of the iron content in the media (Fig S3, S1 Table, S2 Table). Moreover, transcription of Agr-regulated genes [50] was significantly downregulated in the *hemB* mutant (Fig. S3), a characteristic of *S. aureus* SCVs [35,48,51]. We found considerable overlap in the genes upregulated in iron-limited media in the *hemB* mutant and in WT *S. aureus*, where 41 of the 46 genes upregulated in the SCV were also upregulated in the WT (41 of the 46 genes upregulated in the *hemB* SCV) (Fig 2C). In both WT *S. aureus* and the *hemB* mutant the iron storage protein ferritin (*ftnA*) had lower transcript levels while several Fur-regulated iron acquisition system genes had higher transcript levels in iron limited media (Fig 2A and 2B). Indeed, transcript levels of the high-affinity heme iron acquisition system, Isd, and the siderophore acquisition systems (Hts, Sir, Fhu, and Sst) were also significantly higher in both WT *S. aureus* and the *hemB* mutant when the bacteria were grown under iron deplete conditions (Fig 2A and 2B). Interestingly, iron starvation was found to differentially upregulate an additional 54 genes in WT *S. aureus*, including the *sbn* operon required for biosynthesis of staphyloferrin B (SB) (Fig 2C). Under the culture conditions and used here, transcription of the *sbn* operon was below our limit of detection in the *hemB* mutant for either condition (S1 Table). Furthermore, only 6 of the 50 total downregulated genes in iron-limited media were shared among WT and *hemB S. aureus* (Fig 2D); these genes included *ahpC*, encoding an alkyl hydroperoxide reductase and *fur* (S2 Table). Among the genes found to be downregulated by iron depletion in the WT only (and not in the *hemB* mutant) were the *pur* operon genes, involved in purine biosynthesis, and the gene encoding catalase, *katA*. Taken together, our analysis led to the conclusion that both WT *S. aureus* and the *hemB* mutant sense and respond to iron starvation by upregulating expression of iron acquisition systems.

**Figure 2.**
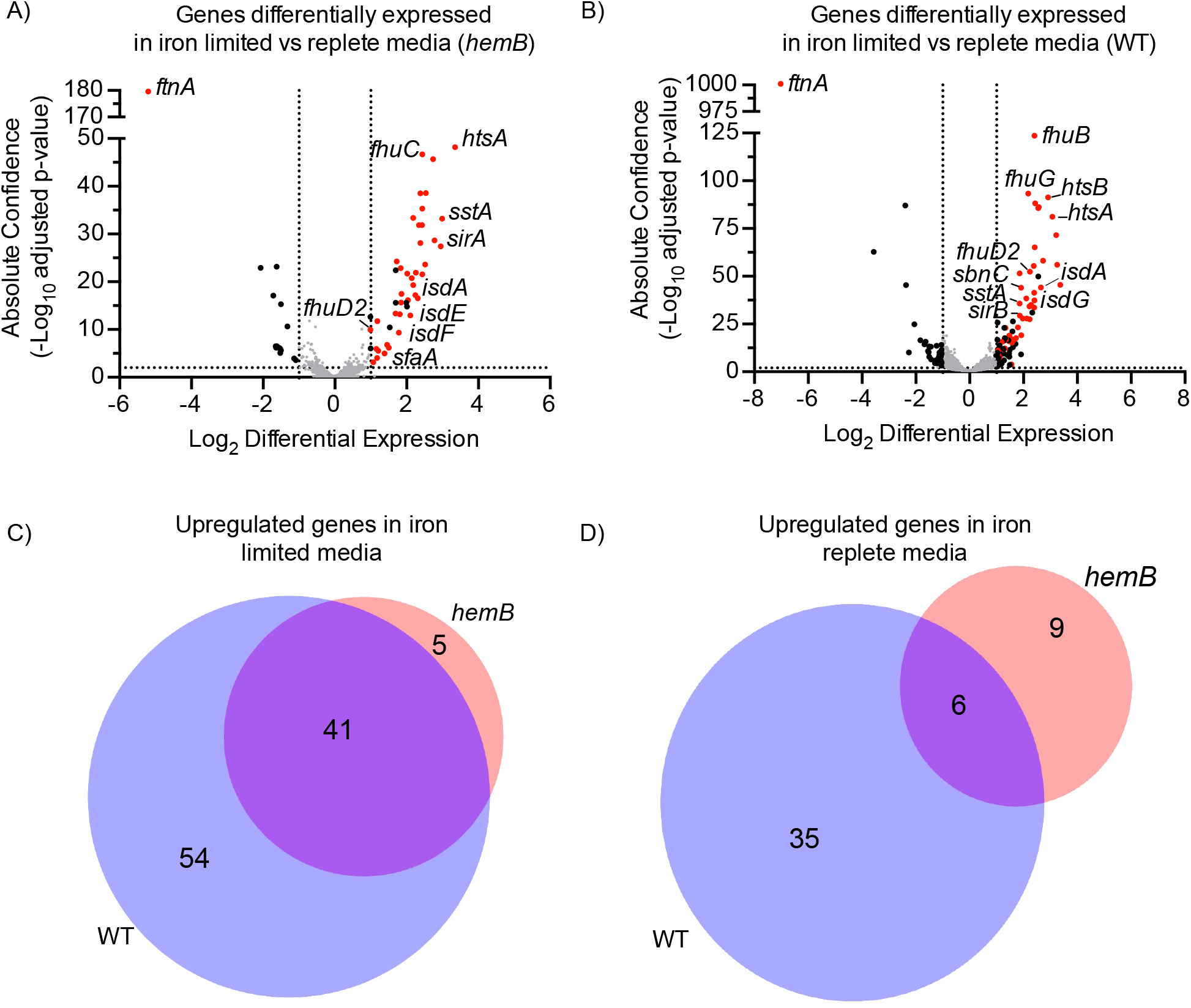
Analysis of RNA expression in response to iron starvation by *S. aureus* WT and *hemB*. Volcano plots of genes that were differentially expressed in iron-limited vs iron-replete media by either **(A)** the *S. aureus hemB* SCV or **(B)** WT *S. aureus*. Significantly up- and down-regulated genes (absolute confidence > 2 and |log_2_ differential expression| > 1) that are Fur-regulated are colored red and remaining significantly regulated genes are colored black. **(C)** Comparison of the number of significantly upregulated or **(D)** significantly downregulated genes in iron limited media by WT and *hemB S. aureus*. RAW reads for RNAseq data can be found in S1 Table, and lists of significantly up- and down-regulated genes in all conditions can be found in S2 Table.

### Prioritization of endogenous staphyloferrin B over staphyloferrin A for iron acquisition by the *S. aureus hemB* SCV

Given that our RNA-seq data revealed differences in gene expression of staphyloferrin biosynthesis genes between WT *S. aureus* and the *hemB* SCV, we investigated the utilization of endogenous siderophores by the *S. aureus hemB* mutant. As discussed above and shown in Fig 1, the *S. aureus hemB* SCV is debilitated for growth under conditions of iron starvation, suggesting that one or both of its endogenous high affinity siderophore-dependent iron acquisition pathways (i.e. staphyloferrin A [SA] and staphyloferrin B [SB]) are defective. To characterize this further, we generated mutants in the *hemB* SCV background that lack either the SA or SB biosynthesis loci, *sfa* or *sbn*, respectively. Growth analyses revealed that the SA biosynthesis-deficient (-SA) *hemB* mutant grew akin to parental *hemB* whereas SB biosynthesis-deficient (-SB) *hemB* bacteria had a significant growth defect in the iron-restricted growth medium (+hTf) (Fig 3A). Given the RNA-seq analyses demonstrating that, at least at mid log phase, *hemB S. aureus* did not display appreciable transcription of the *sbn* operon we sought an alternative means to confirm use of the SB by *hemB* bacteria. To this end staphyloferrin uptake mutants were generated in the *hemB* background that lacked either or both of the dedicated, high-affinity SA and SB receptors, HtsA [52, 53] and SirA [54], respectively. Corroborating the findings above, these analyses demonstrated that under iron deplete conditions only inactivation of *sirA*, which renders the bacteria unable to utilize SB (-SB), significantly impaired growth relative to parental *hemB S. aureus* (Fig 3A). In contrast, inactivation of the *hts* operon, required for SA uptake (-SA) [55, 56], was without effect in the *hemB* background (Fig 3A). Staphyloferrin utilization was shown important only under iron restriction, as *hemB* mutants deficient for either SA or SB biosynthesis and uptake grew as well as their *hemB* isogenic parent strain under iron replete conditions (Fig 3B). Taken together, these data provide compelling evidence that *hemB S. aureus* relies predominantly on utilization of endogenous SB, and not SA, to support a minimal level of growth when iron is scarce and demonstrate that, at least at some point during iron-restricted growth, *hemB S. aureus* must express the *sbn* operon.

**Figure 3.**
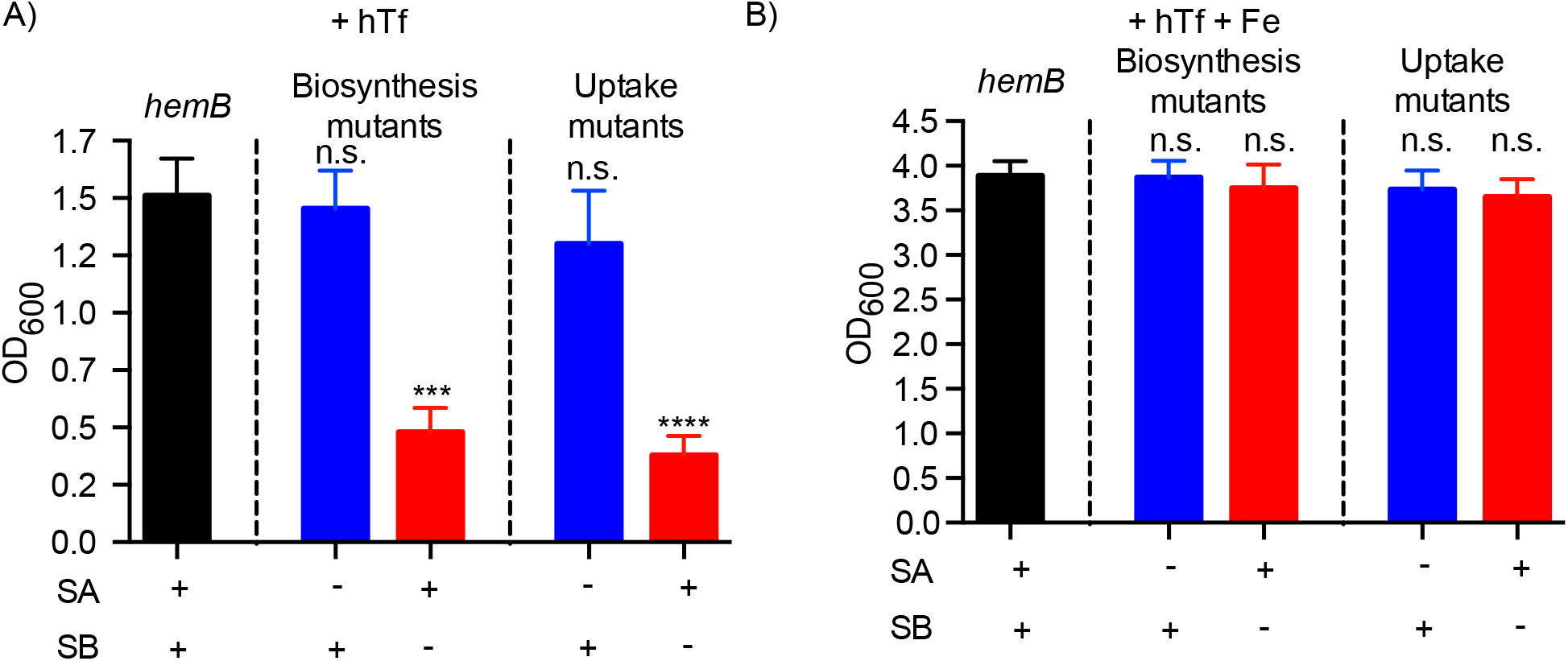
Staphyloferrin B-mediated iron acquisition is required for optimal iron-starved growth of *hemB* SCV. Growth of *hemB S. aureus* (black), or *hemB S. aureus* combined with mutation of SA (blue) or SB (red) iron acquisition systems. SA and SB biosynthetic and uptake mutations are indicated. Bacteria were cultured for 24 h in TMS supplemented with 0.4 µM hemin and **(A)** 1.5 µM human transferrin (+hTf), or **(B)** 1.5 µM human transferrin and 10 µM ferric ammonium sulfate (+hTf+Fe). Data are plotted as mean ± SEM, nine biological replicates from three independent experiments. \*\*\**p* ≤ 0.001, **** *p* ≤ 0.0001, one-way ANOVA with Tukey’s multiple comparisons.

### In a *hemB* SCV, staphyloferrins are dispensable for growth *in vivo*

To determine whether the *S. aureus hemB* mutant requires endogenous SA and/or SB *in vivo*, a murine model of systemic infection was used. Mice were infected with *hemB S. aureus* and a *hemB* mutant defective for staphyloferrin biosynthesis (-SA -SB). This analysis revealed that weight loss of the animals and bacterial counts in the kidneys and liver were similar for the parental *hemB* strain and the staphyloferrin deficient *hemB* mutant (Fig S4). These observations reveal that, *in vivo*, loss of staphyloferrin-dependent iron acquisition is without effect in *hemB S. aureus*, suggesting these bacteria utilize another source of iron, such as heme, to meet their iron requirements.

### The *hemB* mutant displays an exogenous SA utilization deficiency

Considering our finding that the *S. aureus hemB* mutant prioritizes utilization of endogenous SB over SA to support a minimal level of growth in iron-limited media, we next investigated whether the *hemB* mutant can utilize SA when provided exogenously. To this end, growth experiments were performed with the staphyloferrin-deficient SCV and non-SCV *S. aureus* in TMS with hemin and hTf as described above, however, purified SA was also added to the medium. Both the *hemB* SCV (Fig 4A) and non-SCV *S. aureus* deficient for staphyloferrin biosynthesis (-SA -SB) (Fig S5) failed to proliferate in iron deplete media. When added to TMS medium containing hTf, the purified SA supported SA/SB-deficient non-SCV *S. aureus* to fully restore growth. This observation confirmed that the purified SA was a functional siderophore (Fig S5). In contrast, growth of the staphyloferrin deficient *hemB* mutant remained attenuated for growth in iron-restricted medium supplemented with SA, though the purified SA did enable some growth (Fig 4A). Remarkably, the maximal growth achieved by *hemB S. aureus* in iron-limited media through limited utilization of either purified SA or endogenously synthesized SB was virtually identical (Fig 3A and 4A). Given that the *hemB* mutant was limited in its ability to utilize exogenous SA, we next considered whether the SA receptor HtsA is differentially expressed by the SCV. In agreement with our RNA-seq data, western blot analysis of HtsA expression revealed that *hemB S. aureus* express HtsA in an iron dependent manner akin to WT *S. aureus* and the complemented *hemB* mutant (Fig 4B). Taken together, these data demonstrate that the *hemB* SCV has defective, but not completely inoperative, SA utilization.

**Figure 4.**
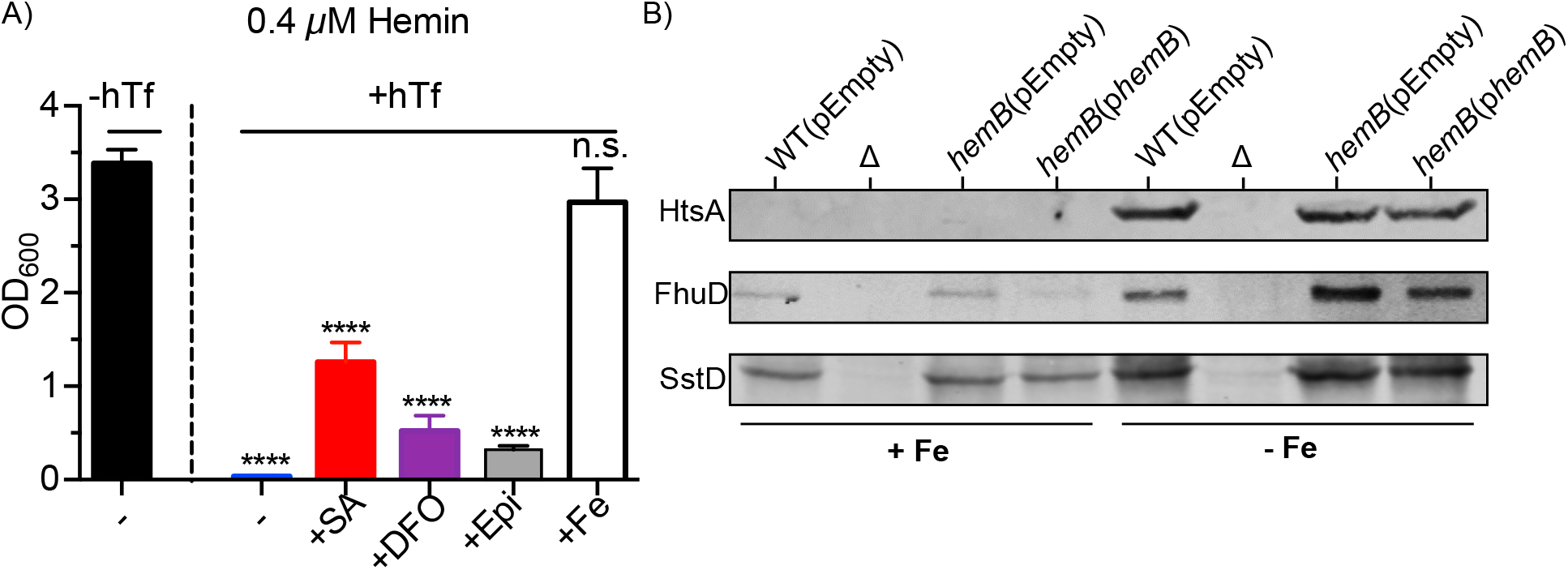
Siderophore utilization by *S. aureus hemB* is defective. Growth of the staphyloferrin deficient *hemB sfa sbn* mutant cultured for 24 h in TMS supplemented with 0.4 µM hemin. Medium contains either no human transferrin (-hTf) or 5 µM human transferrin (+hTf). Either 100 µM of siderophore (staphyloferrin A (+SA), deferoxamine mesylate (+DFO), or epinephrine (+Epi)), or 30 µM ferric ammonium sulfate (+Fe) was added to the iron-restricted medium. Data are plotted as mean ± SEM, nine biological replicates from three independent experiments. **** *p* ≤ 0.0001, one-way ANOVA with Tukey’s multiple comparisons. **(B)** Western blots for detection of HtsA (31 kDa), FhuD2 (34 kDa), or SstD (38 kDa) expression by WT(pEmpty), *hemB*(pEmpty), or *hemB*(p*hemB*). A negative control (Δ), either a *S. aureus htsA*, *fhuD2*, or *sstD* mutant, was used in each respective Western blot. Whole cell lysates were prepared from bacteria cultured for 24 h in TMS supplemented with 0.4 µM hemin and either 30 µM ferric ammonium sulfate (+Fe) or 1.5 µM human transferrin (-Fe).

### The *S. aureus hemB* mutant has deficient xenosiderophore utilization

We next examined whether exogenous provision of xenosiderophores could support growth of the *S. aureus hemB* mutant in iron-deplete media. The clinically approved drug Desferal (the mesylate salt of deferoxamine; DFO) and the stress hormone epinephrine have been shown to act as hydroxamate- and catechol-type siderophores, respectively, that support the growth of *S. aureus* in iron-restricted environments [56, 57]. Therefore, growth experiments were performed with staphyloferrin deficient SCV and non-SCV *S. aureus* as previously described, where either DFO or epinephrine were added to Tris minimal succinate (TMS) medium supplemented with hemin and hTf. As was observed with the exogenous provision of SA, growth the staphyloferrin deficient *hemB* mutant was not restored by the provision of xenosiderophores, though again some limited growth was observed (Fig 4A). In contrast, provision of DFO or epinephrine rescued growth of non-SCV *S. aureus* deficient for staphyloferrin biosynthesis, indicating the xenosiderophores were indeed functional as agents of iron delivery (Fig S5). Given that xenosiderophore utilization was impaired in the *S. aureus hemB* mutant, we examined whether the receptors for DFO or epinephrine, FhuD2 [58] and SstD [56], respectively, were differentially expressed by the *hemB* SCV by western blot. This analysis revealed that FhuD2 and SstD were both expressed in an iron-dependent manner by *hemB S. aureus*, comparably to WT *S. aureus* and the complemented *hemB* mutant (Fig 4B), which corroborated our RNA-seq data. These results demonstrate that the *S. aureus hemB* SCV has deficient xenosiderophore utilization despite expression of hydroxamate- and catechol-siderophore acquisition systems at both the RNA (Fig 2) and protein (Fig 4) levels.

### Examination of the intracellular ATP levels of the *hemB* SCV

HemB is required for endogenous heme biosynthesis and the terminal oxidases in the electron transport chain of *S. aureus* utilize heme as co-factor [40]. Therefore, we hypothesized that that the inability of *hemB* bacteria to utilize siderophores for growth could be due to disruption of electron transport and ATP production. Indeed, previous reports have suggested that SCV bacteria produced less ATP than WT bacteria [33, 42]. To test this notion, we measured the intracellular ATP of WT and *hemB S. aureus* grown in TSB at various timepoints. Remarkably, intracellular ATP increased in both WT and the *hemB* mutant during exponential growth phase for each respective strain (Fig 5A and 5B). Due to the prolonged lag phase of the *hemB* mutant, the observed increase in intracellular ATP was delayed by ∼4 hours as compared to WT *S. aureus* (Fig 5A and 5B). We next examined the influence of Fe on the ability of WT and *hemB S. aureus* to generate ATP. Here the bacteria were grown in iron restricted (+hTf) or iron replete (+hTf + Fe) TMS supplemented with hemin. These data revealed that *hemB* bacteria, under conditions of Fe limitation, were able to generate ATP akin to WT however, the onset of ATP production was again delayed in *hemB S. aureus* as compared to WT (Fig 5C-D and S6). Furthermore, comparison of ATP production between WT and *hemB S. aureus* in Fe replete and restricted conditions, revealed that Fe supplementation augmented ATP production. Once again, peak ATP production was delayed in the *hemB* background as compared to WT (Fig 5E and 5F). Taken together, these data demonstrate that the ability of *hemB* bacteria to generate ATP is not markedly different from WT and therefore cannot explain the observed siderophore utilization defects that significantly curtail iron-restricted growth.

**Figure 5.**
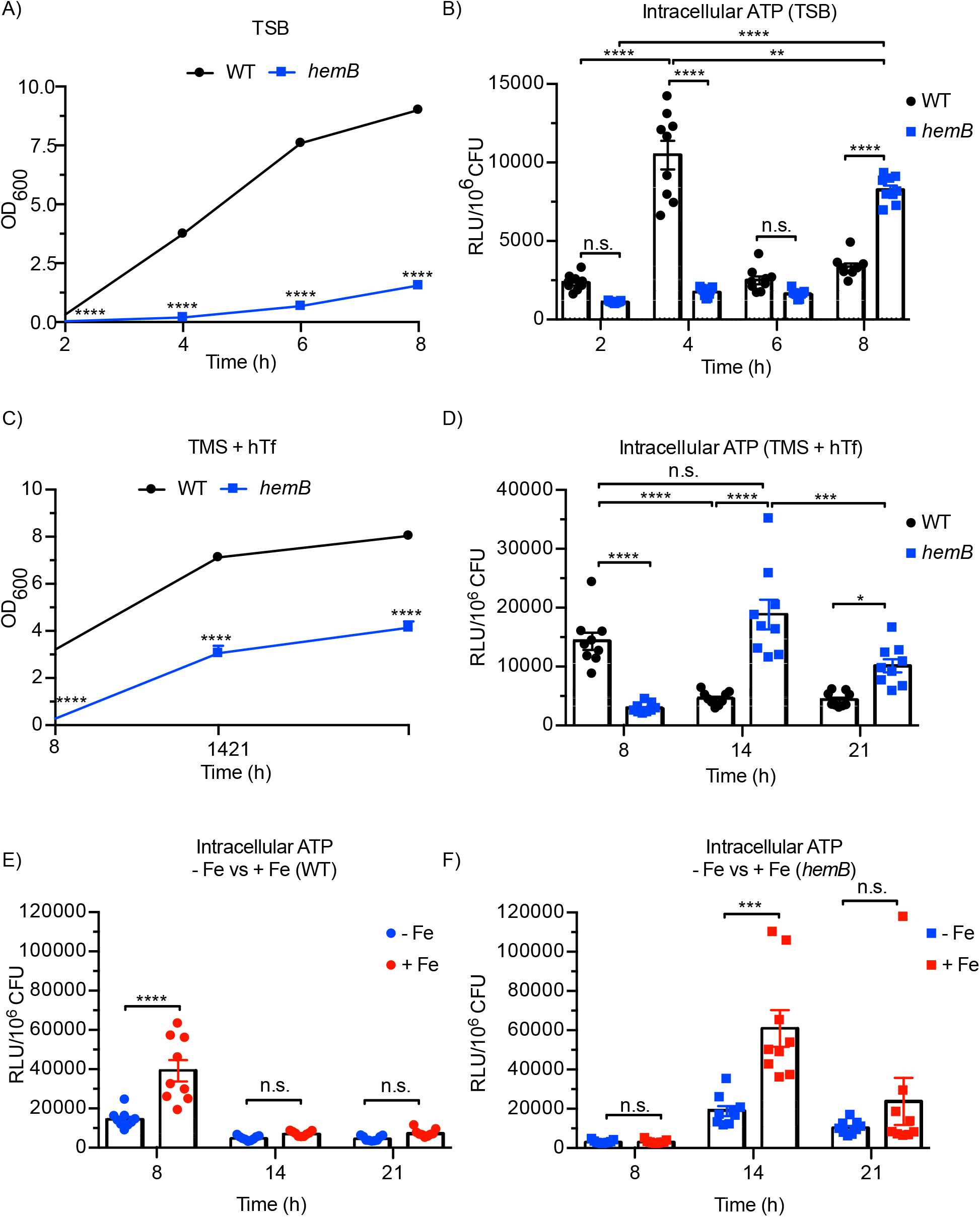
Intracellular ATP of a *S. aureus hemB* mutant. Growth **(A)** and intracellular ATP levels of WT and *hemB S. aureus* grown in TSB. Growth **(C)** and intracellular ATP levels **(D)** of WT and *hemB S. aureus* grown in TMS supplemented with 0.4 µM hemin and 1.5 µM human transferrin (+hTf). Intracellular ATP levels of WT *S. aureus* **(E)** or the *hemB* mutant **(F)** grown in TMS supplemented 0.4 µM hemin, 1.5 µM hTf, and either 0 µM ferric ammonium sulfate (-Fe) or 10 µM ferrous ammonium sulfate (+Fe). Data are plotted as mean ± SEM, nine biological replicates from three independent experiments. * *p*<0.05, ** *p*<0.01, *** *p*<0.001, **** *p*<0.0001, two- way ANOVA with Dunnett’s multiple comparisons **(A, C)** one-way ANOVA with Tukey’s post test **(B, D-F)**.

### The *hemB* mutant demonstrates niche-specific utilization of DFO *in vivo*

Previously, it was demonstrated that administration of DFO to *S. aureus*-infected mice augmented virulence in a systemic model of infection [59]. To examine whether DFO could enhance *S. aureus hemB* growth *in vivo*, systemic infections were performed using mice that were administered DFO intraperitoneally. In this model, infections were terminated at 48 h post-infection because WT infected mice having received DFO met humane endpoint criteria and had lost a significant amount of body weight as compared to mice not having received DFO (Fig 6A). In contrast, mice infected with the *S. aureus hemB* mutant retained significantly more weight irrespective of DFO administration, indicating that *hemB* bacteria were attenuated *in vivo* (Fig 6A). At this time the bacterial burden in the hearts, kidneys, and livers of infected mice was also determined. This analysis revealed that WT *S. aureus* was better able to infect the heart and kidneys of infected animals as compared to *hemB* bacteria. Moreover, administration of DFO augmented WT *S. aureus* USA300 growth in every organ analyzed (Fig 6B-D). In contrast, DFO failed to augment growth of *hemB S. aureus* in the liver and heart of infected mice (Fig 6B and 6D). Remarkably, DFO administration led to a modest, yet significant, increase in *hemB* bacteria in the kidneys of infected mice (Fig 6C). Conceivably, within the kidney *hemB* bacteria access a pool of heme that in part complements the auxotrophy and enables DFO utilization in this niche. Nevertheless, these data demonstrate that while DFO dramatically increases the virulence of WT *S. aureus in vivo*, DFO is largely without effect on the *hemB* SCV mutant. These *in vivo* data agree with data presented in Fig 4, illustrating that DFO utilization by the *hemB* mutant, *in vitro*, is severely attenuated.

**Figure 6.**
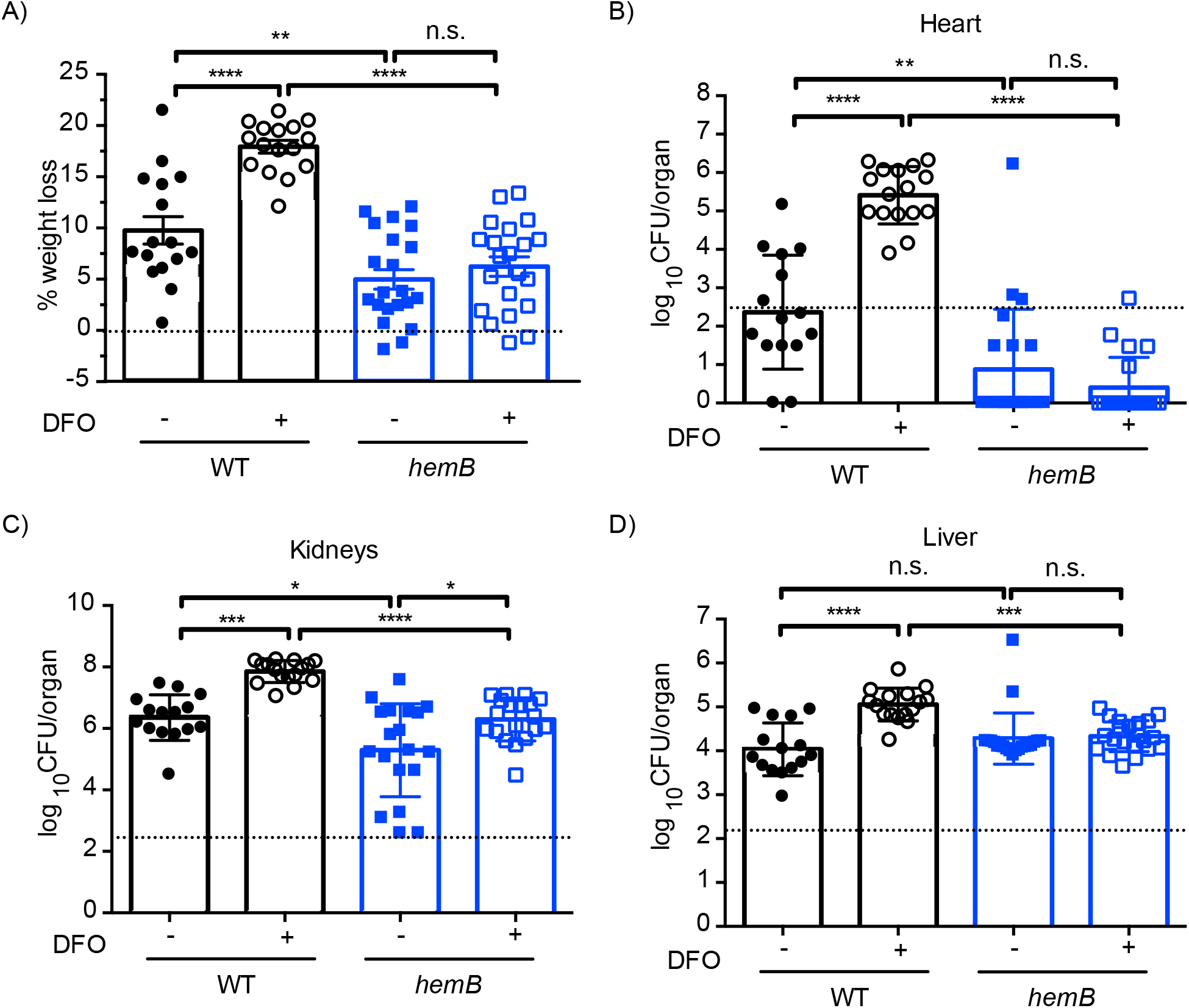
Niche-specific utilization of DFO by *S. aureus hemB in vivo* in a murine model of systemic infection. Mice were inoculated with *S. aureus* USA300 WT or *hemB* and received either DFO or vehicle control (see Materials and Methods). 48 h post infection, mice were sacrificed and the **(A)** percent weight loss and bacterial burden of the heart **(B)**, kidneys **(C)**, or liver **(D)** was determined. Limit of accurate detection is represented as a dashed line. Data are plotted as mean ± SEM, at least 15 animals per group. * *p* ≤ 0.05, ** *p* ≤ 0.01, *** *p* ≤ 0.001, **** *p* ≤ 0.0001, one-way ANOVA with Tukey’s post test.

### Provision of additional hemin enhances DFO utilization by the *hemB* mutant

Spurred by our finding that DFO augmented *hemB* mutant growth exclusively in the murine kidney, we posited that the availability of heme affects DFO utilization. To examine whether provision of additional heme enhances hydroxamate-type siderophore utilization by the *hemB* mutant, growth experiments were performed in the presence of DFO where the TMS growth medium was supplemented with either 0.4 µM or 2 µM hemin. Previously we found the addition of 2 mM hemin could increase but not fully complement the growth rate, colony size, and pigmentation of the *S. aureus hemB* mutant (Fig S1). Remarkably, DFO augmented growth of the *hemB* mutant in iron-restricted medium when the culture was supplemented with 2 µM hemin (Fig 7A). Importantly, that the bacteria fail to demonstrate enhanced growth in the presence of 2 mM hemin (c.f. 0.4 mM hemin) without DFO indicates that the additional hemin on its own does not provide sufficient iron to a *hemB* SCV to overcome hTf-dependent growth restriction (Fig 7A). To determine whether the ability of *hemB S. aureus* to use DFO in the presence of additional heme was specific to catechol type siderophores we repeated the same growth experiments however exogenous SA and epinephrine were provided. Here we utilized a *hemB* mutant also lacking the *sfa* and *sbn* operons so the bacteria could not utilize any endogenously synthesized siderophore. In the presence of 2 mM hemin, DFO was again utilized by staphyloferrin-deficient *hemB* bacteria, however the bacteria were also now able to utilize SA (Fig 7B). Indeed, in the presence of SA and 2 mM hemin the staphyloferrin-deficient *hemB* mutant was able to grow to a significantly higher optical density as compared to conditions with only 0.4 mM hemin (see Fig 7B and 5A, respectively). In contrast, provision of additional hemin did not enable epinephrine utilization by staphyloferrin-deficient *hemB S. aureus*, indicating some siderophore utilization defects persist (Fig 7B). Importantly, epinephrine was readily used by non-SCV siderophore-deficient *S. aureus* to overcome hTfn iron restriction demonstrating the functionality of epinephrine as a siderophore (Fig S5). Given the effect of hemin on siderophore utilization we also sought to measure the effects of hemin on ATP production. This analysis demonstrated that *hemB* and WT *S. aureus* produce a similar amount of ATP during exponential growth irrespective of hemin concentration. However, when grown in a medium containing 2 mM hemin, both strains generated more ATP as compared to the 0.4 mM hemin condition (Fig 7C). Taken together, these data demonstrate that siderophore utilization by *hemB S. aureus* can be influenced by extracellular hemin availability. Moreover, these data reveal that *hemB S. aureus* utilize heme as a nutrient to support siderophore transport and utilization to overcome iron restriction as opposed to as an iron source alone.

**Figure 7.**
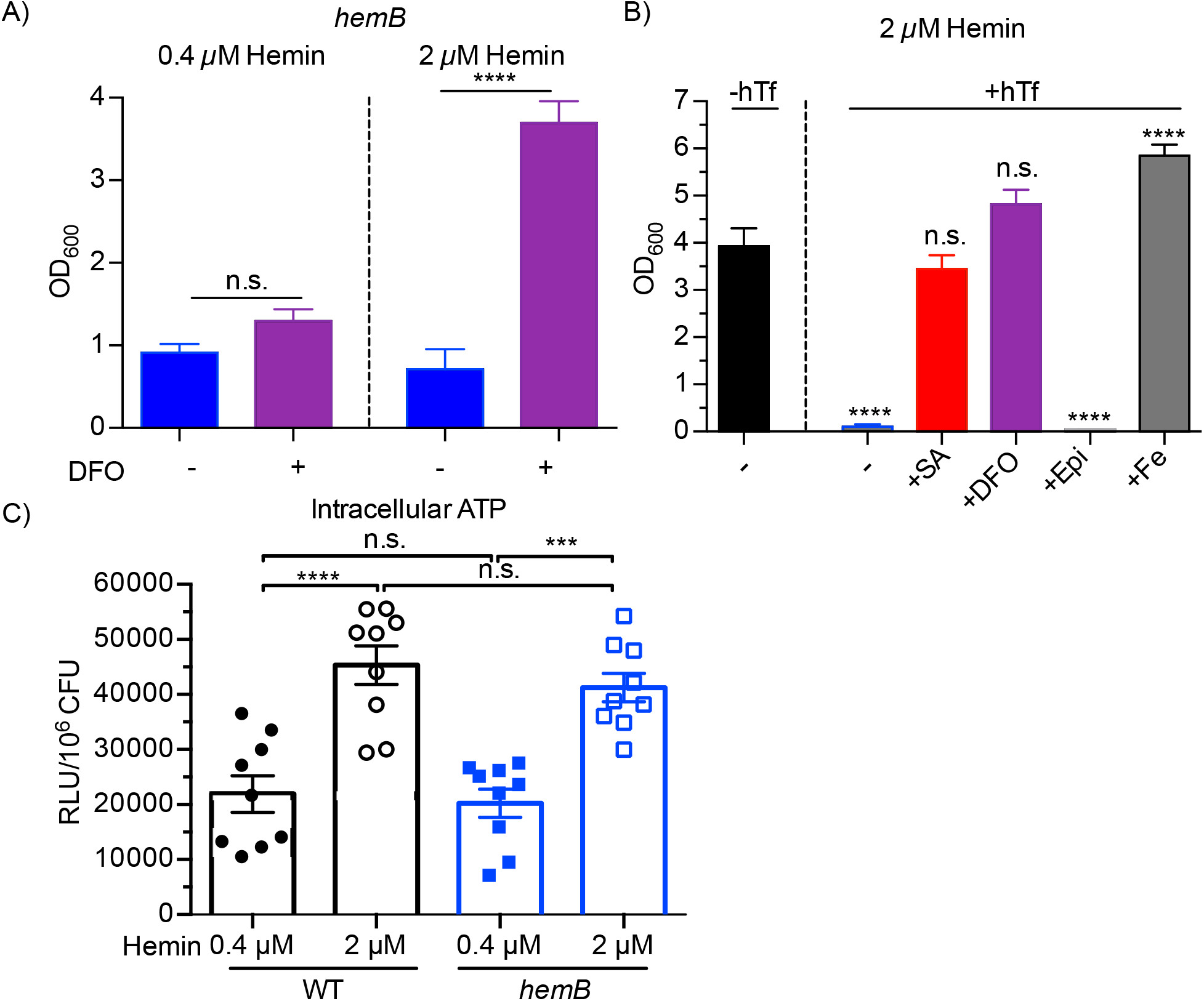
Siderophore utilization by *S. aureus hemB* is enhanced by additional hemin. (A) Growth of the *hemB* mutant cultured at 37°C for 24 h in iron-restricted TMS supplemented with either 0.4 µM or 2 µM hemin, as indicated. Medium contains 5 µM hTf to restrict iron and either 0 µM or 100 µM DFO to assess utilization of the hydroxamate-type siderophore. Data are plotted as mean ± SEM, at least five biological replicates. **** *p* ≤ 0.0001, one-way ANOVA with Tukey’s multiple comparisons. **(B)** Growth of the staphyloferrin deficient *hemB sfa sbn* mutant cultured at 37°C for 24 h in TMS supplemented with 2 µM hemin. Medium contains either no human transferrin (-hTf) or 5 µM human transferrin (+hTf). Either 100 µM of siderophore (staphyloferrin A (+SA), deferoxamine mesylate (+DFO), or epinephrine (+Epi)), or 30 µM ferric ammonium sulfate (+Fe) was added to the iron-restricted medium. **(C)** Intracellular ATP levels of WT and *hemB S. aureus* grown to exponential phase in TMS supplemented with either 0.4 µM or 2 µM hemin. Data are plotted as mean ± SEM, nine biological replicates from three independent experiments. *** *p* ≤ 0.001, **** *p* ≤ 0.0001, one-way ANOVA with Tukey’s multiple comparisons.

## Discussion

Small colony variants of *S. aureus* occur naturally during infection. These variants are often characterized by their slow growth, inability to hemolyze red blood cells, and their enhanced antimicrobial resistance [34,36,60,61]. That *S. aureus* SCVs cause chronic persistent infections necessitates the bacteria acquire essential nutrients, such as iron, within the host. During infection iron is scarce due to host driven mechanisms of nutrient sequestration that function to deprive pathogens and curtail replication [18, 21]. Consequently, *S. aureus* has evolved several specialized high-affinity iron acquisition systems, powered by ATP hydrolysis, to overcome host iron sequestration [24]. Here, we sought to determine the iron acquisition strategies employed by a *S. aureus* SCV. Since clinical SCV isolates readily revert to the wild-type state [62, 63], we generated a stable *S. aureus* SCV by mutating the *hemB* gene. Indeed, *hemB* mutants of *S. aureus* have been utilized as an archetypal SCV in numerous studies [42,47,64–68]. The *hemB* gene encodes a delta-aminolevulinic acid dehydratase (aka porphobilinogen synthase) that acts in the classical pathway for heme biosynthesis which commences with the conversion of 5- aminolevulinic acid to uroporphyrinogen III [69]. The importance of heme as a prosthetic group is underscored by its conservation in virtually all forms of life and explicably, disruption of the *hemB* gene in *S. aureus*, has pleiotropic effects [42]. How disruption of endogenous heme biosynthesis affects iron acquisition by *S. aureus* has to our knowledge not been explored. Our experiments reveal that *hemB S. aureus*, under conditions of iron restriction, displays impaired siderophore utilization despite the bacteria retaining the ability to express several high affinity iron acquisition systems (see Fig 2 and Fig 4). Indeed, our work demonstrated that *hemB S. aureus* could sustain only limited growth in iron deplete media through diminished utilization of endogenous staphyloferrin B (SB), or even through exogenous addition of siderophores such as staphyloferrin A (SA), deferoxamine B (Desferal) or epinephrine (see Fig 3, 4 and 7). In contrast, these iron chelates fully correct the iron restricted growth of a siderophore-deficient non-SCV *S. aureus*. Given *hemB S. aureus* express the machinery needed for siderophore acquisition (e.g. SirA, HtsA, FhuD1, FhuD2 and SstD) we also considered whether defects in siderophore utilization could be attributed to aberrant ATP production as previous work has indicated that SCV bacteria have a reduced capacity to synthesize ATP [70, 71]. To this end we measured intracellular ATP content of WT and *hemB* bacteria cultured in TMS and hemin as described in methods. This analysis revealed that *hemB S. aureus* can produce ATP at similar levels, as compared to WT, however the timing of peak ATP production is delayed in *hemB* bacteria ostensibly due to the drastic differences in growth kinetics (see Fig 5). Taken together our data indicate that an inability to synthesize ATP alone cannot explain the siderophore utilization defects by the *S. aureus* SCV. Conceivably, the inability of *hemB* bacteria to utilize siderophores is due to defects in iron extraction from siderophores once transported into the cell [72–75] and future work will aim to test this possibility.

That we utilized a *hemB* mutant, which has previously been shown to be auxotrophic for hemin [42], necessitated the use of a minimal amount of hemin to permit bacterial growth in the medium employed here. Careful consideration was given to the concentration of hemin used in our experiments, as excess hemin could conceivably complement the *hemB* mutant and altogether eliminate the SCV phenotype. Importantly, the low concentration of hemin employed here (0.4 mM) permitted proliferation of *hemB S. aureus* without correcting the SCV growth defect as compared to WT bacteria (see Fig 1) indicating the effects of *hemB* disruption were maintained. Furthermore, RNA-seq reaffirmed that the effects of *hemB* disruption were not complemented upon provision of a minimal amount of hemin, as significant differences between the transcriptome for *hemB* and wild-type *S. aureus* were still observed under the conditions analyzed (Fig S3), including downregulation of Agr-regulated genes, a characteristic trait ascribed to SCVs of *S. aureus* [49]. In addition to complementing the *hemB* mutant auxotrophy, the requirement for hemin in our experiments was complicated by the fact that hemin itself could act as an iron source. Indeed, *S. aureus* can utilize nanomolar concentrations of host-derived heme as a preferred iron source for growth [31, 76]. Despite this, the growth data presented here (e.g. Fig 7) indicate that under the iron-restricted conditions we employed, even as much as 2 µM hemin, in the presence of hTf to sequester free iron, does not provide a sufficient amount of iron to allow *S. aureus* to overcome hTf-dependent metal sequestration. Furthermore, the observation that 2 µM hemin cannot provide sufficient iron to support *hemB* mutant growth demonstrates that in the presence of siderophore the bacteria utilize the hemin provided as a nutrient that enables siderophore utilization. The mechanism by which this occurs is not immediately clear however our data reveal that differences in the ability to generate ATP cannot account for the observed phenotype. Siderophores as low molecular weight iron scavenging molecules bind ferric iron (Fe^3+^) with extraordinary affinity while displaying very low affinity for ferrous iron (Fe^2+^). Interestingly, liberation of Fe^3+^ from siderophore complexes within the bacteria cell requires Fe reduction. In *S. aureus* two reductases have been identified IruO and NtrA that are required for the utilization of the DFO and SA, respectively [75]. In a *hemB S. aureus* mutant the activity of the IruO and NtrA or whether other reductive mechanisms could operate has not been characterized. It is tempting to speculate that a heme-containing protein may be required for siderophore reduction in *hemB S. aureus*, the function of which is influenced by exogenous heme. Indeed, in *Escherichia coli*, the flavohaemoglobin protein Hmp displays ferrisiderophore reductase activity [77] and whether a heme-containing protein also functions in siderophore reduction in *hemB S. aureus* requires further study.

WT *S. aureus* can synthesize two siderophores, SA and SB, which can support growth under iron-restricted conditions. Interestingly, our data show that SA biosynthesis is dispensable for *hemB* mutant growth in the presence of hTf however, SB biosynthesis is not. While our bacterial growth data convincingly demonstrate that SB biosynthesis and transport are required for the optimal iron-restricted growth of the *hemB* SCV, the RNA-seq data regarding the lack of detectable levels of *sbn* operon genes is intriguing. We attribute this to sampling at an early stage of log phase growth when, presumably, heme is at its highest concentration in the medium. We have previously shown that heme serves as a signal, through the SbnI protein, that *S. aureus* uses to downregulate expression of *sbn* genes to prioritize heme utilization [78]. The apparent prioritization of siderophore utilization (SB >> SA) may, in part, be attributable to important metabolic differences between wild-type *S. aureus* and the SCV. For instance, previous work has shown that the tricarboxylic acid (TCA) cycle is downregulated in *hemB S. aureus* [47, 79] and, in wild-type *S. aureus*, citrate from the TCA cycle is required for SA biosynthesis [79]. Interestingly, our RNA-seq data reveal that in the *hemB* mutant the citrate synthase gene *citZ* is not transcriptionally downregulated, however, the gene encoding the aconitase hydratase *acnA*, also part of the TCA cycle, is (see S2 Table). Therefore, it stands to reason that reduced citrate availability in a *hemB* mutant, because of perturbation of the TCA cycle, results in a significant deficiency in synthesis of SA leading to very limited growth (see Fig 3A). Notably, many previous studies have complemented *hemB* mutant phenotypes with exogenous heme in rich growth media such as TSB, which would provide enough free iron in addition to the heme to allow robust growth of the *hemB* SCV. Indeed, we note in this study that when provided as little as 0.4 µM heme, free iron added to the minimal medium robustly complements the growth defect of the *hemB* mutant. Of note, this form of iron would not require reductase enzymes for iron release inside the cell as is the case for iron complexed to siderophores. *In vivo*, however, bacteria are rarely presented with this form of iron as it is complexed to host iron-binding proteins, components of nutritional immunity imparted on microbes by the host.

It is well accepted that siderophore production by *S. aureus* contributes to bacterial fitness during infection [56, 79], however, siderophore expression can be heterogenous and influenced by the niche occupied by the bacteria [80]. That accessibility to iron in the host is indeed limited, despite the ability of WT *S. aureus* to produce siderophores, is evidenced by the observation that intraperitoneal injection of DFO significantly enhances bacterial replication and virulence *in vivo* [59], as it does for other pathogens such as *Yersinia enterocolitica* [81]. The *in vivo* role or significance of SA and SB in the context of the *hemB* mutation in *S. aureus* is, at present, unclear given that elimination of the *sfa* and *sbn* loci does not further attenuate the bacterial load in organs infected with the *hemB* SCV. This would be concordant with our *in vitro* data demonstrating that *hemB S. aureus* poorly utilize siderophores when iron restricted (see Fig 3 and 4). Conceivably, heme, the preferred iron source of *S. aureus* [82], is the predominant source of iron for the *hemB S. aureus* during infection however the contribution of heme acquisition systems, such as the Isd pathway, to SCV growth remain to be explored. It is noteworthy that SCVs will arise *in vivo* as part of a mixed population of *S. aureus* comprised of WT and SCVs. Previous work has shown that the auxotrophy of some SCVs can be cross-complemented through the metabolite production by other *S. aureus* bacteria, or even by the microbiome, as has been aptly demonstrated by Hammer *et al*. [83]. However, in the case of iron acquisition specifically, we have shown here that the iron-restricted growth defect of the *hemB* mutant cannot be adequately complemented by many siderophore-iron complexes; note that ferric ammonium sulfate (or ferric chloride, data not shown) can complement the iron-restricted growth of a *hemB* SCV (see Fig 1, 4 and 7) but, as stated above, ‘free’ iron rarely exists in the host as it is sequestered by host iron-binding proteins [18]. Ostensibly, because the host represents a hostile, iron-restricted niche, it is unlikely that *hemB* SCVs would utilize siderophores produced by other bacteria unless also presented with a rich source of heme. Consequently, *hemB* bacteria would be expected to remain as a reservoir of antibiotic resistant, fitness-defective bacteria until such a time as they genetically revert to the WT state. In this regard, work is currently underway in the laboratory to examine iron-restricted growth characteristics of other types of SCVs, such as menadione or thymidine auxotrophic SCVs.

Consistent with previous reports [83–85] we find that the *hemB S. aureus* SCV is attenuated relative to WT *S. aureus* using a systemic murine model of infection. Furthermore, our experiments evaluating *in vivo* utilization of the iron chelator desferrioxamine (DFO, or Desferal) further emphasized the profound differences between the virulence of WT *S. aureus* and the *hemB* mutant. Indeed, growth of WT *S. aureus* was significantly enhanced in every organ analyzed upon intraperitoneal injection of DFO. In contrast, the *hemB* mutant was afforded no DFO-mediated growth enhancement in either the heart or liver of infected animals. Remarkably, however, DFO administration conferred a modest yet significant growth enhancement to the *hemB* mutant in kidneys of infected animals. Our *in vitro* data revealed that *hemB* mutant bacteria can utilize DFO as an iron source provided that their nutritional requirement for heme is met. This aligns with our suggestion that within the kidneys of infected animals SCVs can access heme, likely because of high bacterial numbers that invoke tissue damage and heme release. Previous work using a rabbit endocarditis model of infection demonstrated that *hemB S. aureus* could proliferate similar to WT in the kidneys of infected rabbits [86]. Presumably, in this instance, *hemB* bacteria access heme through tissue damage and/or through the extreme sensitivity of rabbit red blood cells to lysis by alpha-toxin despite reduced *hla* expression by SCV bacteria [87].

In summary, the work presented here demonstrates that a *hemB* SCV in *S. aureus* displays inherent defects in siderophore-dependent growth under iron-restricted conditions, and that these defects are reliant on heme concentration. Despite this, organ specific differences in heme availability may influence siderophore utilization and the subsequent growth of *hemB* bacteria. Understanding how SCVs acquire essential nutrients during infection will facilitate the identification of novel drug targets that, if perturbed, could help treat chronic *S. aureus* SCV infection. Indeed, perturbation of heme acquisition may represent a strategy whereby the ability of *hemB* bacteria to acquire iron during infection can be prevented.

## Materials and Methods

### Bacterial strains, plasmids and growth media

All bacterial strains and plasmids used in this study are listed in Table 1. *S. aureus* USA300 LAC cured of its endogenous antibiotic resistance plasmid served as the WT strain for this study. *Escherichia coli* DH5α was used for cloning purposes and was cultured in Luria-Bertani broth (LB; Difco). *S. aureus* strains were grown in either Tryptic Soy broth (TSB; Wisent) or Tris Minimal Succinate (TMS) [88]. All media and solutions were prepared using water purified with a Milli-Q water filtration system (EMD Millipore, Billerica, MA). Solid media were supplemented with 1.5% (w/v) Bacto agar (Difco). The antibiotics spectinomycin (300 µg/mL), chloramphenicol (12 µg/mL), tetracycline (4 µg/mL), and kanamycin (50 µg/mL) were added to media as required for *S. aureus*; ampicillin (100 µg/mL) was added where required for *E. coli*. Bacteria were incubated at 37°C, with constant shaking (225 rpm) where required, unless indicated otherwise.

**Table 1.**
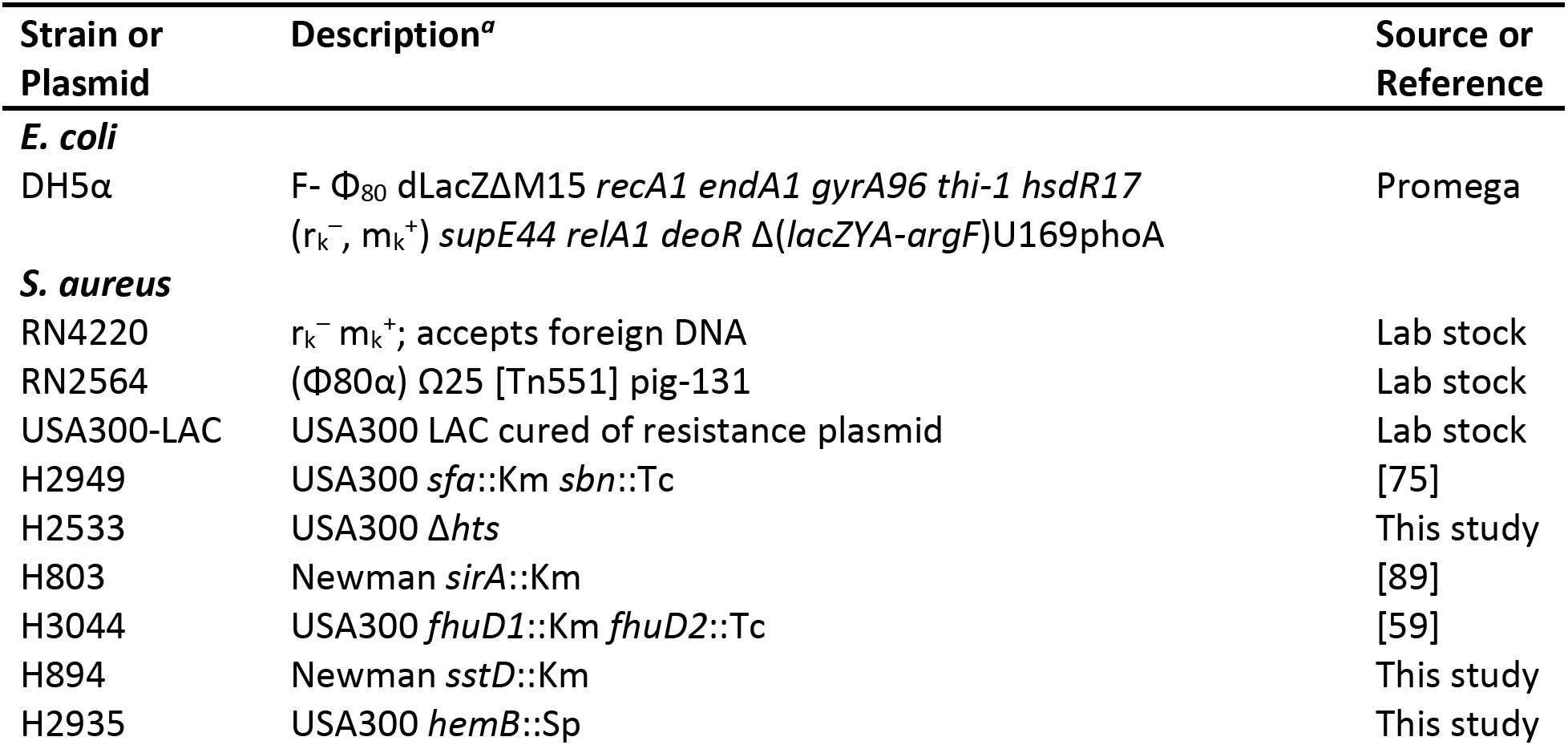
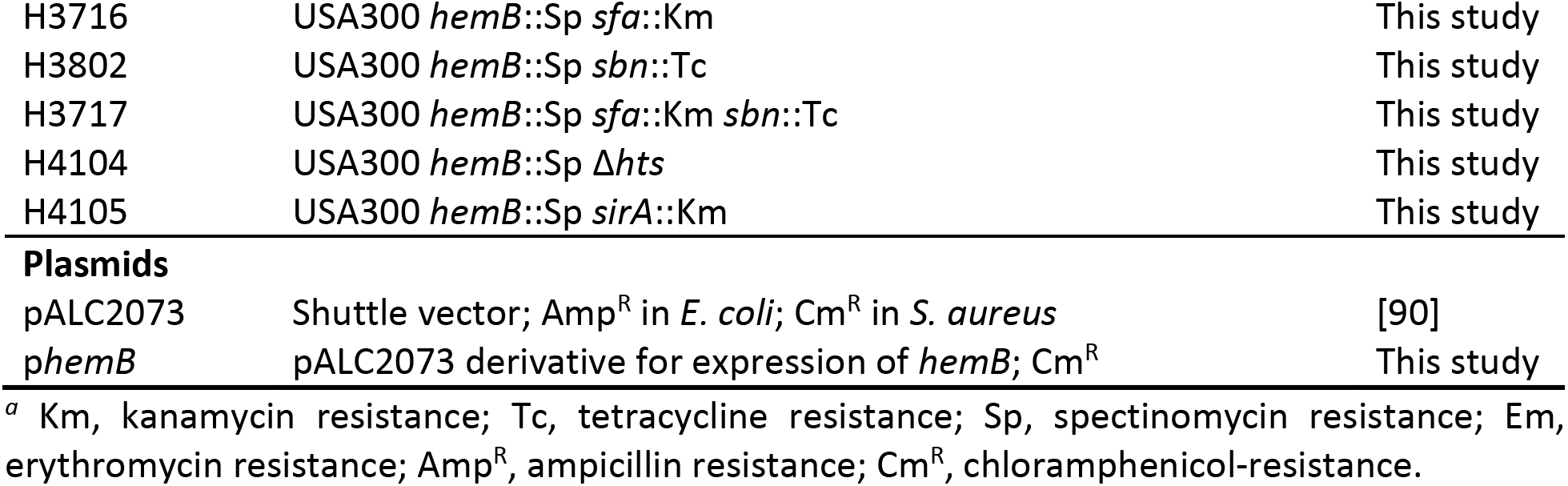
Bacterial strains and plasmids employed in this study.

| Strain or Plasmid | Description<sup>a</sup> | Source or Reference |
| --- | --- | --- |
| ***E. coli*** |  |  |
| DH5α | F- $\Phi_{80}$ dLacZΔM15 *recA1 endA1 gyrA96 thi-1 hsdR17* ($r_k^-$, $m_k^+$) *supE44 relA1 deoR* $\Delta$(*lacZYA-argF*)U169phoA | Promega |
| ***S. aureus*** |  |  |
| RN4220 | $r_k^-$ $m_k^+$; accepts foreign DNA | Lab stock |
| RN2564 | (Φ80α) Ω25 [Tn551] *pig*-131 | Lab stock |
| USA300-LAC | USA300 LAC cured of resistance plasmid | Lab stock |
| H2949 | USA300 *sfa::Km sbn::Tc* | [75] |
| H2533 | USA300 Δ*hts* | This study |
| H803 | Newman *sirA::Km* | [89] |
| H3044 | USA300 *fhuD1::Km fhuD2::Tc* | [59] |
| H894 | Newman *sstD::Km* | This study |
| H2935 | USA300 *hemB::Sp* | This study |
| H3716 | USA300 *hemB*::Sp *sfa*::Km | This study |
| H3802 | USA300 *hemB*::Sp *sbn*::Tc | This study |
| H3717 | USA300 *hemB*::Sp *sfa*::Km *sbn*::Tc | This study |
| H4104 | USA300 *hemB*::Sp Δ*hts* | This study |
| H4105 | USA300 *hemB*::Sp *sirA*::Km | This study |
| Plasmids |  |  |
| pALC2073 | Shuttle vector; Amp<sup>R</sup> in *E. coli*; Cm<sup>R</sup> in *S. aureus* | [90] |
| *phemB* | pALC2073 derivative for expression of *hemB*; Cm<sup>R</sup> | This study |

<sup>a</sup> Km, kanamycin resistance; Tc, tetracycline resistance; Sp, spectinomycin resistance; Em, erythromycin resistance; Amp<sup>R</sup>, ampicillin resistance; Cm<sup>R</sup>, chloramphenicol-resistance.

### Mutant construction and complementation

The *hemB* mutant was obtained by interrupting the gene with a spectinomycin resistance cassette and integration into the chromosome using published procedures [91]. For mobilizing the *hemB*::Sp and *sirA*::Km mutations into various genetic backgrounds, phage transduction was performed using standard techniques. Briefly, phage lysate was prepared from the donor strain using Φ80α (prepared from RN2564). Recipient strains were infected, transductants were selected using appropriate antibiotics, and insertions were confirmed by PCR. Primers used in this study are summarized in Table 2. For complementation of the *hemB* mutation, the full-length gene was amplified using the *hemB*-F-SacI and *hemB*-R-KpnI primer pair (Table 2), ligated into pALC2073 and transformed into *E. coli*. The p*hemB* complementation plasmid was passaged through *S. aureus* strain RN4220 before introduction of p*hemB* into the strain of interest; an equivalent procedure was performed for introduction of the empty vector pALC2073 into various strains. All chromosomal mutations were confirmed by sequencing PCR amplified products from across the relevant regions of the genome.

**Table 2.**
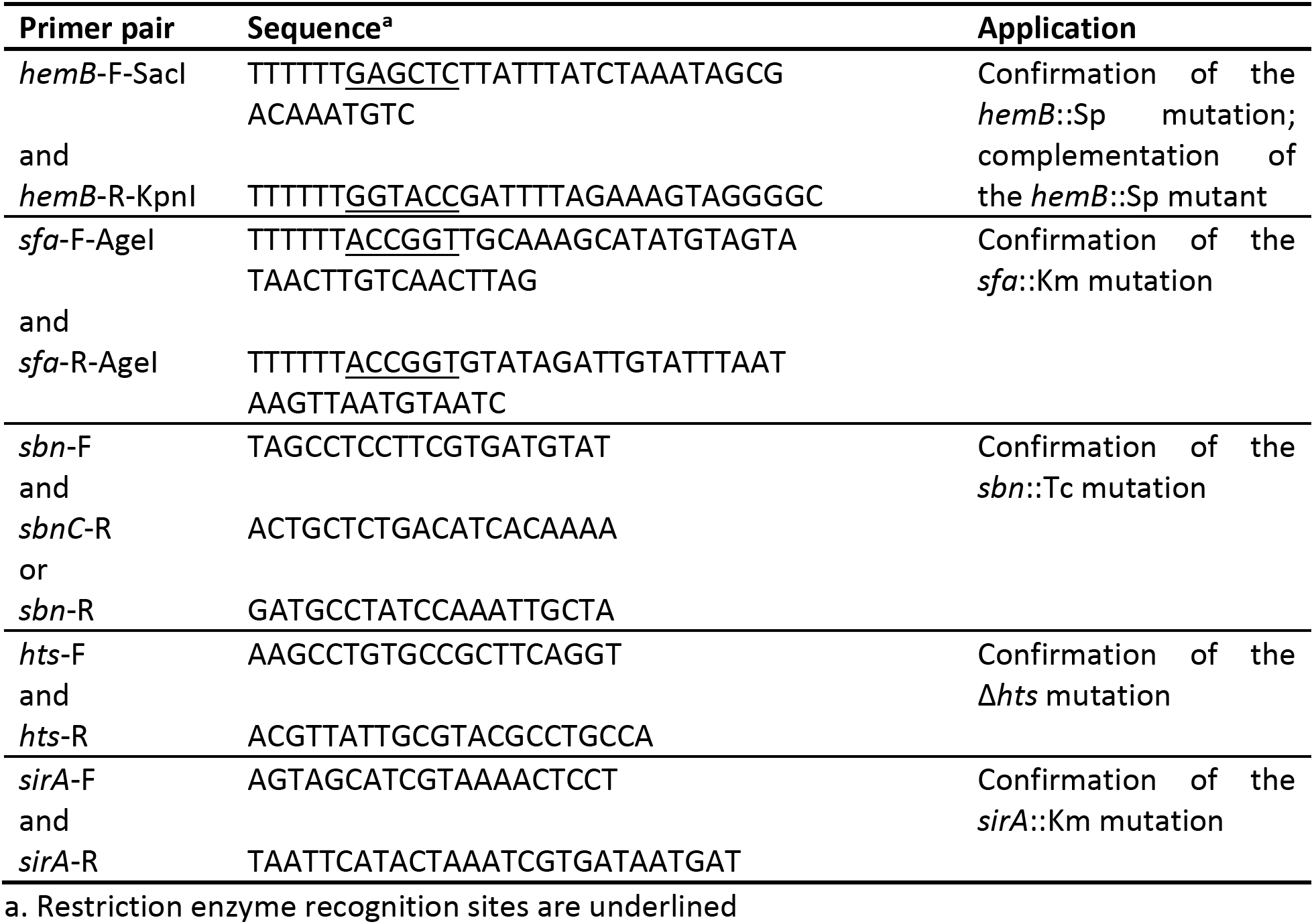
Oligonucleotides employed in this study.

### Bacterial growth assays

To assess growth of *S. aureus* in TSB or TMS, single isolated colonies were picked from TSA plates and were inoculated into TSB. Overnight cultures of bacteria were washed twice with sterile saline and normalized to an OD_600_ of 0.1. TSB or TMS inoculated with diluted *S. aureus* to give a starting OD_600_ of 0.001. Bacterial growth was assessed by measuring the OD_600_ after 24 h or every 3 h over the course of 24 h for growth curves. Subsequent growth assays in TMS were similarly performed, but overnight cultures were grown in TMS and TMS was supplemented with hemin, as indicated. Hemin stocks were prepared using bovine hemin (Sigma), as previously described [78].

### Phenotypic assessment on TSA plates

Phenotypic assessment of the *hemB* mutant, relative to WT USA300 (carrying the empty vector pALC2073) and the *hemB*-complemented strain (containing p*hemB*) was performed on TSA plates supplemented with either 0 μM or 2 μM hemin. Single isolated colonies were picked from TSA plates and used to inoculate TSB. Overnight cultures of bacteria were washed twice with sterile saline and normalized to an OD_600_ of 0.1. This suspension was serially diluted 10-fold through to 10^-4^ and 10 μL of each dilution was drop plated on TSA plates with either 0 μM or 2 μM hemin. Plates were incubated at 37°C for 48 h and representative images were taken. To assess toxin production, a single isolated colonies of WT *S. aureus*, the *hemB* mutant, and the complemented *hemB* mutant were patched to TSA plates supplemented with 5% (v/v) human blood. Plates were incubated at 37°C for 24 h and representative images were taken.

### Assessment of siderophore utilization

Human apo-Tf (hTf; Sigma) was used to restrict iron availability in TMS. Iron restriction in media was alleviated by the addition of ferrous ammonium sulfate (FAS; VWR), which was a source of free Fe^3+^. Purified staphyloferrin A (SA; Indus Biosciences Private Limited), Desferal^®^ (deferoxamine mesylate, DFO; Hospira, obtained from the London Health Sciences Centre), or epinephrine (Sigma) were added to iron restricted media to assess siderophore utilization. The concentration of hTf, ferric ammonium sulfate (FAS), and siderophore added was specified for each experiment, as detailed in Results or Figure Legends.

### Preparation of cells for RNA extraction

Single colonies of WT *S. aureus* and the *hemB* mutant were inoculated into TMS supplemented with 0.4 µM hemin and grown overnight. Overnight cultures were subcultured to give a starting OD_600_ equivalent of 0.001 in TMS supplemented with 0.4 µM hemin and 1.5 µM hTf. Cultures were grown until mid-exponential phase, at which point the cultures were spiked with either 0 or 10 µM ferrous ammonium sulfate. Cultures were grown for another hour. An equivalent of OD_600_ = 3.0 of cells were collected and RNAprotect Bacteria Reagent (Qiagen) was mixed with the cells as per manufacturer’s instructions. Cell pellets were frozen overnight at -80°C.

### RNA extraction and RNA sequencing

To extract RNA from frozen WT and *hemB S. aureus*, cell pellets were washed with TE buffer (pH 8.0) and re-suspended in a Tris-HCl mixture with 1 mg/mL lysostaphin (pH 7.5) to lyse for 1 h at 37°C. RNA was extracted using an Aurum^TM^ Total RNA Mini Kit (BioRAD #7326820) as per manufacturer’s instructions. TURBO^TM^ DNase (ThermoFisher Scientific) was used to cleave contaminating DNA and RNA was further purified by phenol/chloroform extraction. Concentration and purity of RNA was determined using a Nanodrop. RNA samples were processed by the Microbial Genome Sequencing Center in Pittsburgh, PA. Ribosomal RNA (rRNA) was depleted (Qiagen FastSelect5S/16S/23S) and cDNA libraries were generated (Qiagen Total Stranded RNA). cDNA libraries were sequenced using NextSeq 550 using a 75bp x 8bp x 8bp setup. RNAseq experiments were performed in biological triplicate. Raw sequencing data have been deposited in GEO Omnibus and are available under the record GSEXXXXX.

### Bioinformatic Analyses of RNA sequencing Data

Sequencing reads were mapped to the reference *S. aureus* USA300 FPR3757 genome in Geneious Prime 2020.1.2 (https://www.geneious.com). Expression analysis and comparisons were performed using DESeq2 [92].

### Western blots

To examine expression of *S. aureus* siderophore-binding lipoproteins HtsA, FhuD2, and SstD, Western blot analyses were performed. *S. aureus* strains were grown overnight in TMS supplemented with minimal hemin, overnight cultures were washed twice with 0.9% (w/v) saline, normalized, and used to inoculate TMS supplemented with minimal hemin and either additional Fe^3+^ (in the form of FAS) or hTf to give an initial OD_600_ of 0.001. Cultures were grown for 24 h, normalized to an OD_600_ of 1.0, and pelleted by centrifugation (19,000 × g for 2 min). To prepare cell lysates, bacterial pellets were resuspended in 75 µL lysis buffer with lysostaphin (50 μg lysostaphin resuspended in 1.25 mL of lysis buffer (25 mM Tris-HCl, 50 mM glucose, 150 mM NaCl, 10 mM EDTA; pH 8.0)) and incubated at 37°C for 1 h. After lysis, 25 µL 4 x Laemmli buffer (240 mM Tris-HCl, pH 6.8, 8% (w/v) SDS, 40% (v/v) glycerol, 0.04% (w/v) bromophenol blue) was added and *S. aureus* samples were boiled for 10 min. Proteins were separated by SDS-polyacrylamide gel electrophoresis (SDS-PAGE) on a 12% polyacrylamide gel. After electrophoresis, standard protocols were followed to transfer proteins to a nitrocellulose membrane. The membrane was blocked with 8% (w/v) skim milk for 3 h before addition of primary antibody (rabbit anti-HtsA, rabbit anti-FhuD2, or rabbit anti-SstD antiserum (diluted 1:1000)). Membranes were incubated with primary antibody overnight at 4°C before addition of secondary antibody (donkey anti-rabbit IgG antibody, DyLight 800 conjugated (diluted 1:20,000); Rockland Immunochemicals, Inc., Limerick, PA). An Odyssey CLx Imaging System and LI-COR Image Studio 4.0 software (LI-COR Biosciences) were used to image the membranes.

### Murine model of systemic infection

All animal experiments were performed in compliance with guidelines set out by the Canadian Council on Animal Care. All animal protocols were reviewed and approved by the University of Western Ontario Animal Use Subcommittee, a subcommittee of the University Council on Animal Care. Six-week-old female Balb/c mice (Charles River laboratories) were injected via tail vein with 100 µL of bacterial culture, containing approximately 7 × 10^6^ – 1 × 10^7^ colony-forming units (CFU) of *S. aureus* bacteria. To prepare the bacteria, strains were grown to OD_600_ 2-2.5 in TSB, washed twice with phosphate buffered saline (PBS) and resuspended to OD_600_ 0.2 in PBS, corresponding to a cell density of approximately 7 × 10^7^ – 1 × 10^8^ CFU/mL. Mice were weighed at the time of infection and infections were allowed to proceed for 48 h before animals were euthanized, reweighed, and organs were aseptically harvested in ice-cold PBS + 0.1% (v/v) Triton X-100 (Thermo Fischer Scientific). Extracted organs were homogenised in a Bullet Blender Storm (Next Advance, Troy, NY) using metal beads, serially diluted, and plated on TSA for enumeration of bacterial burden, presented as log_10_ CFU per organ.

For animal experiments involving DFO treatment, 100 µL of a 10 mg/mL solution of DFO (suspended in sterile PBS) were administered intraperitoneally, one dose at the time of bacterial challenge, and a second dose 24 h post-infection. For the no treatment group, 100 µL PBS (vehicle control) was administered intraperitoneally, in parallel to DFO injection. The dose of DFO administered over the course of the two first days of infection was the same as previously employed by Arifin *et. al* [59] and corresponded to approximately 50 mg/kg/day, which is comparable to the weight-adjusted dose recommended for use in humans (Hospira).

### Intracellular ATP assays

To assess intracellular ATP, bacterial growth assays were performed, as described, with a starting OD_600_ of 0.01 in either TSB or TMS. As indicated, TMS was supplemented with 0.4 µM hemin or 2 µM hemin, and either 1.5 µM hTf or 1.5 µM hTf and 10 µM ferrous ammonium sulfate for the iron-deplete and iron-replete conditions, respectively. At each time point, the OD_600_ was measured, samples were plated to enumerate CFU/mL, and intracellular ATP was determined using the BacTiter-Glo assay kit (Promega) following manufacturer’s instructions. Sample luminescence was determined using a Synergy H4 microplate reader (BioTek).

### Statistical analysis

All statistical analyses and graph production were performed using GraphPad Prism software (GraphPad Software, La Jolla, CA).

## Acknowledgements

The authors thank members of the Heinrichs laboratory for critical review of the manuscript.

## Supplementary Information Legends

**S1 Table. RNA-seq data sets.**

**S2 Table. Lists of genes significantly up- or down-regulated by iron.**

**S1 Fig.**
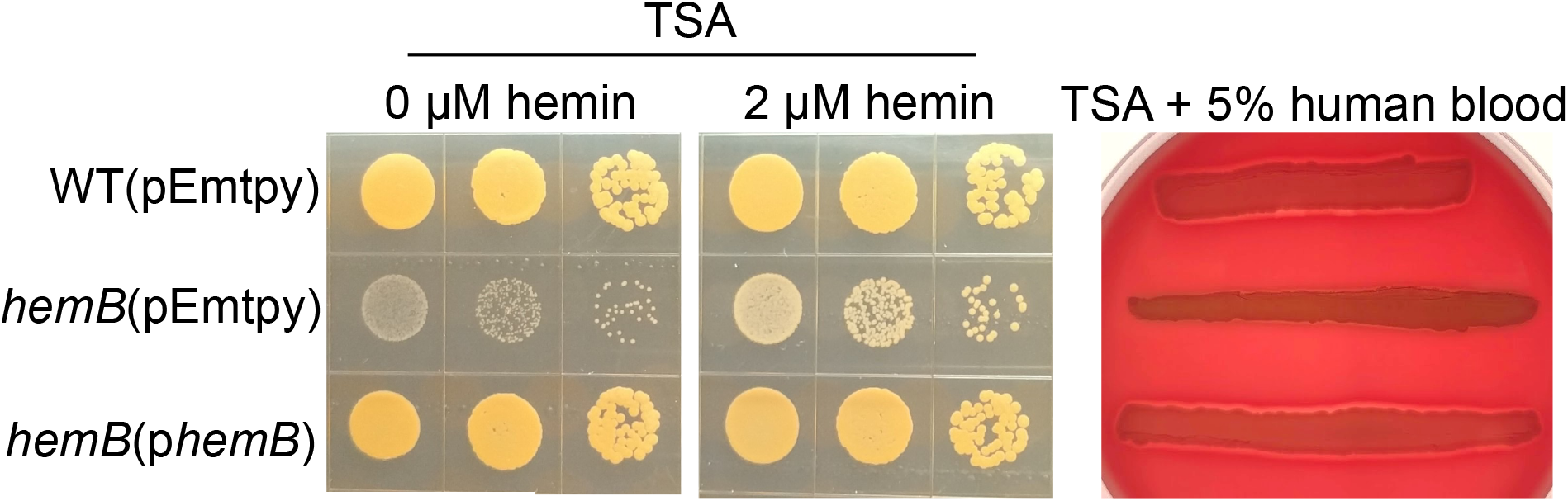
Phenotypic characteristics of the *S. aureus* USA300-LAC *hemB* mutant. Representative images of *S. aureus* USA300 WT(pEmpty), *hemB*(pEmpty), and *hemB*(p*hemB*) grown at 37°C for 48 hr on TSA with or without supplementation of 2 µM hemin or 5 % whole human blood.

**S2 Fig.**
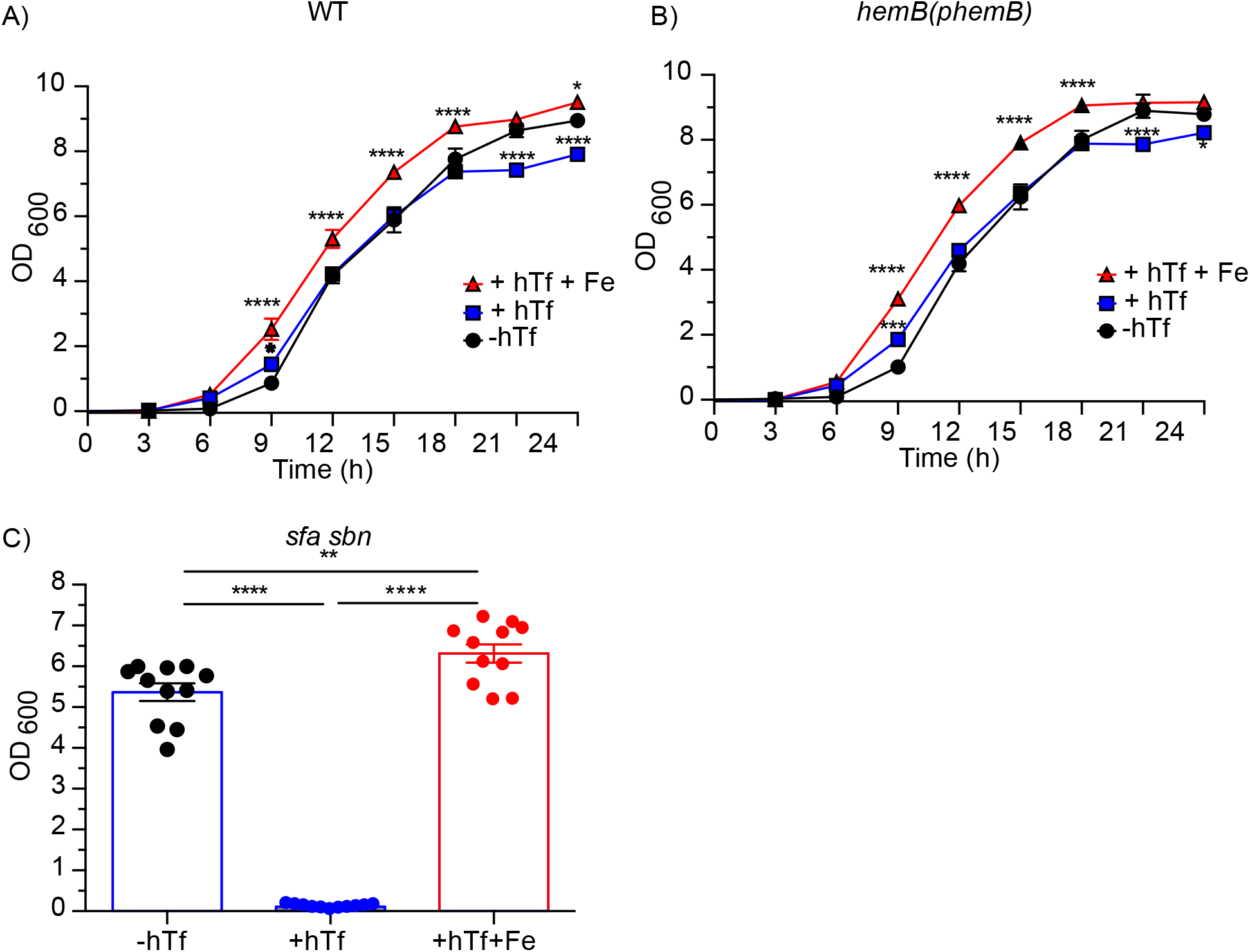
Growth of WT, *hemB*(p*hemB*), and staphyloferrin deficient *S. aureus* in iron-restricted minimal medium. Growth at 37°C over 24 h of **(A)** *S. aureus* USA300 WT or **(B)** *hemB*(p*hemB*) cultured in TMS supplemented with 0.4 µM hemin and either no human transferrin (-hTf), 1.5 µM human transferrin (+hTf), or 1.5 µM human transferrin and 10 µM ferric ammonium sulfate (+hTf+Fe). Data are plotted as mean ± SEM, five biological replicates. \**p* ≤ 0.05, \*\*\**p* ≤ 0.001, **** *p* ≤ 0.0001, two-way ANOVA with Dunnett’s multiple comparisons. **(C)** Growth of *S. aureus sfa/sbn* (-SA -SB) cultured at 37°C for 24 h in TMS supplemented with 0.4 µM hemin and either no human transferrin (-hTf), 1.5 µM human transferrin (+hTf), or 1.5 µM human transferrin and 10 µM ferric ammonium sulfate (+hTf+Fe). Data are plotted as mean ± SEM, 11 biological replicates. \*\**p* ≤ 0.01, \*\*\*\**p* ≤ 0.0001, one-way ANOVA with Tukey’s multiple comparisons.

**S3 Fig.**
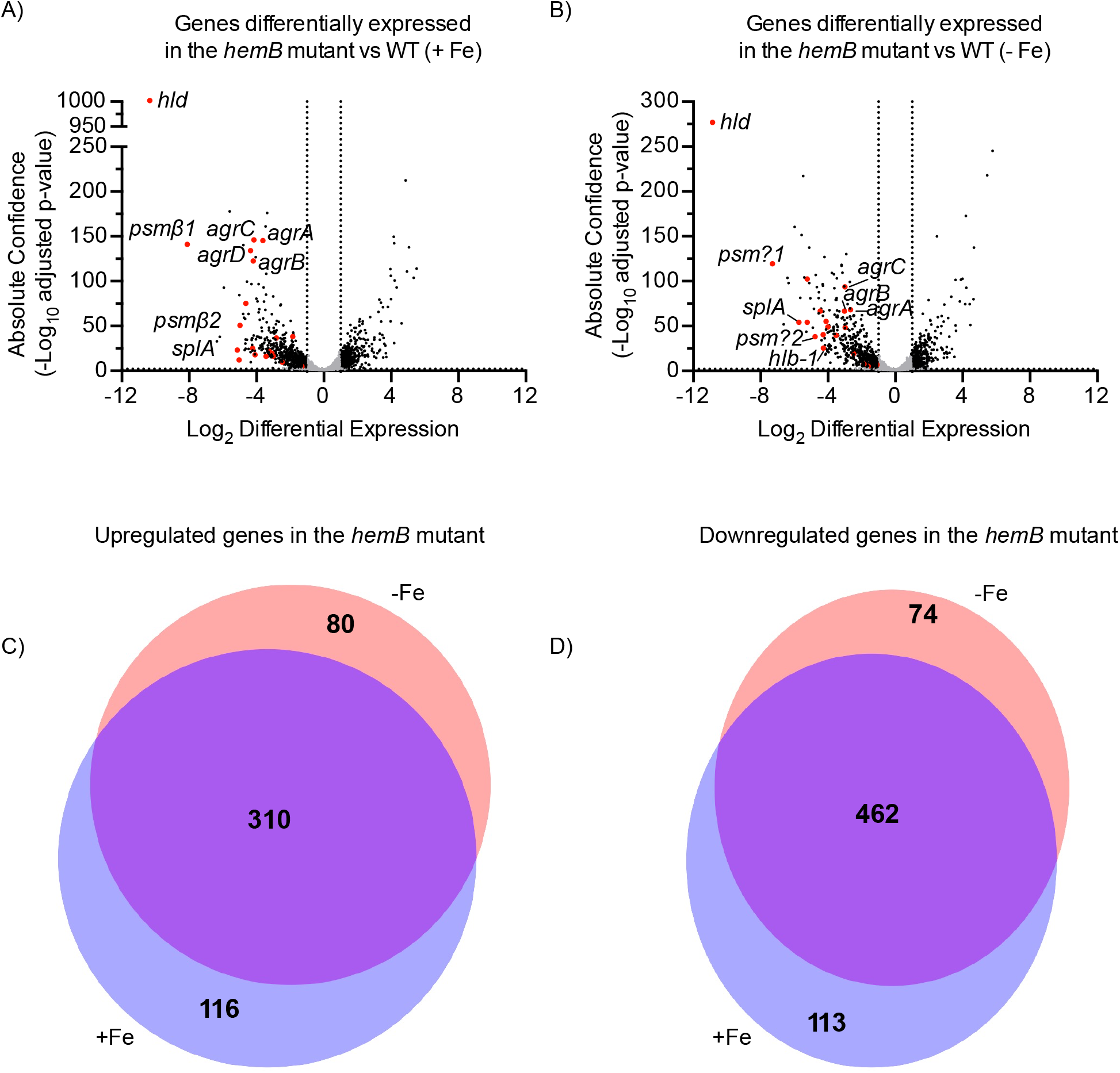
Analysis of RNA expression confirms that *S. aureus hemB*, grown in 0.4 µM, has a transcriptional profile indicative of a classical SCV. Volcano plots of genes that were differentially expressed in the *hemB* SCV as compared to WT *S. aureus* in either **(A)** iron replete media (+ Fe) or **(B)** iron deplete media (- Fe). Significantly up- and down-regulated genes (absolute confidence > 2 and |log_2_ differential expression| > 1) that are Agr-regulated are colored red and remaining significantly regulated genes are colored black. Comparison of the upregulated **(C)** or downregulated **(D)** genes by the *S. aureus hemB* mutant in iron replete and deplete media.

**S4 Fig.**
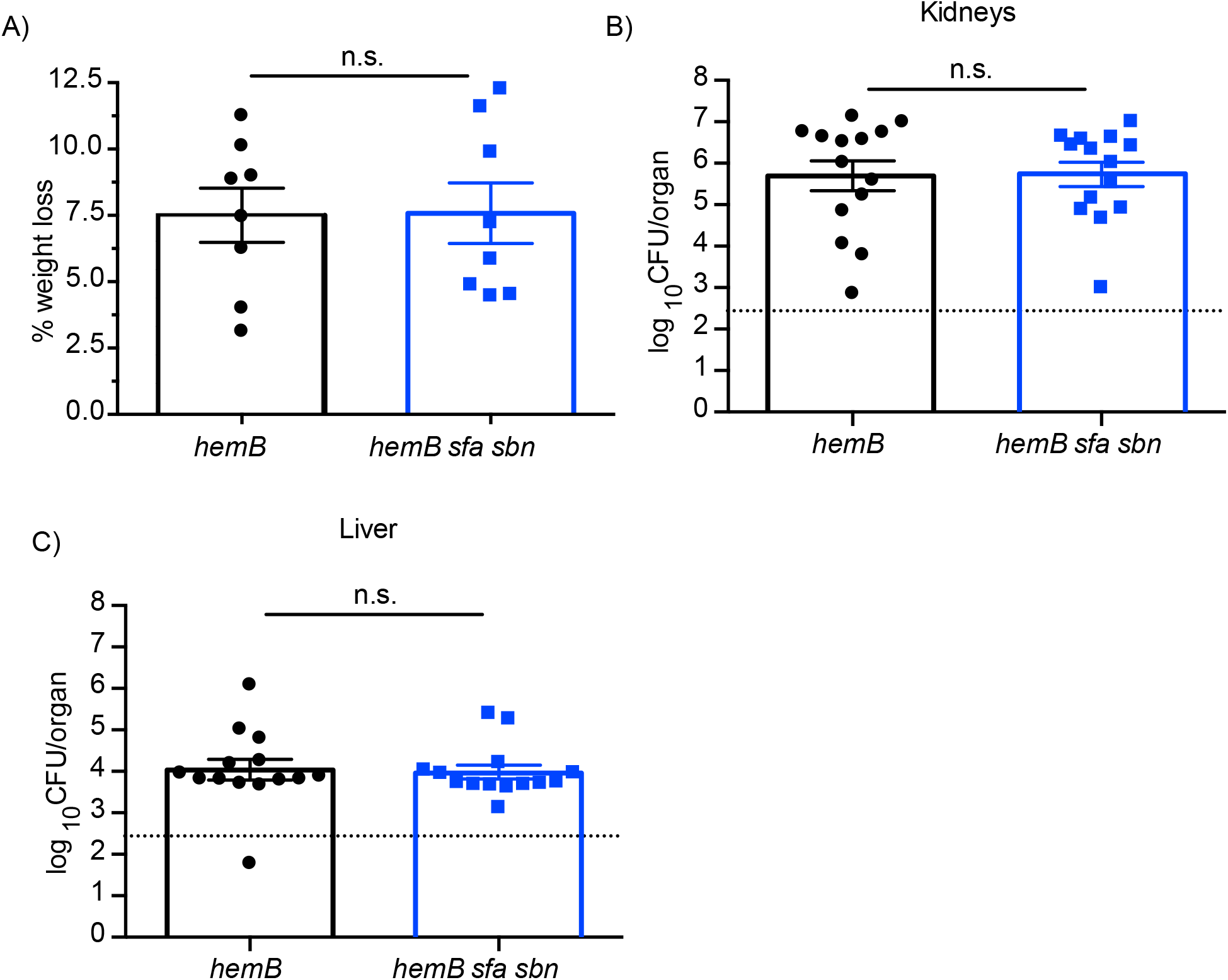
Loss of staphyloferrin expression in a *S. aureus hemB* does not further attenuate bacterial burden in a systemic model of infection. Mice were infected intravenously with *S. aureus hemB* or the staphyloferrin deficient strain *hemB sfa sbn*. 48 h post infection, mice were sacrificed and percent weight loss **(A)** and bacterial burden of the kidneys **(B)** and liver **(C)** was determined. Limit of accurate detection is represented as a dashed line. Data are plotted as mean ± SEM, **(A)** 8 animals per group, **(B-C)** 14 animals per group from two independent experiments. One-way ANOVA with Tukey’s post test.

**S5 Fig.**
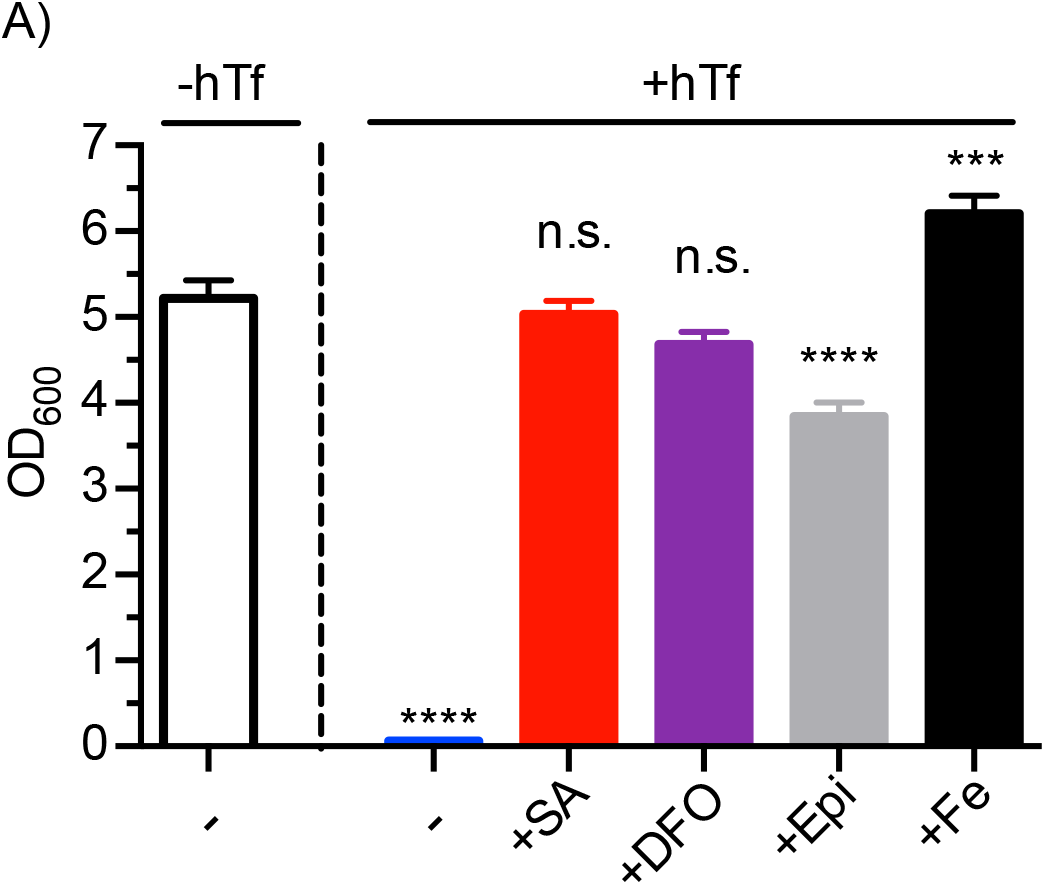
Confirmation that exogenous siderophores function to provide iron to staphyloferrin- deficient *S. aureus*. Growth of the *S. aureus sfa sbn* mutant cultured at 37°C for 24 h in TMS supplemented with 0.4 µM hemin. Medium contains either no human transferrin (-hTf) or 5 µM human transferrin (+hTf). Either 100 µM of siderophore (staphyloferrin A (+SA), deferoxamine mesylate (+DFO), or epinephrine (+Epi)), or 30 µM ferric ammonium sulfate (+Fe) was added to the iron-restricted medium. Data are plotted as mean ± SEM, at least four biological replicates. *** *p* ≤ 0.001, **** *p* ≤ 0.0001, one-way ANOVA with Tukey’s multiple comparisons.

**S6 Fig.**
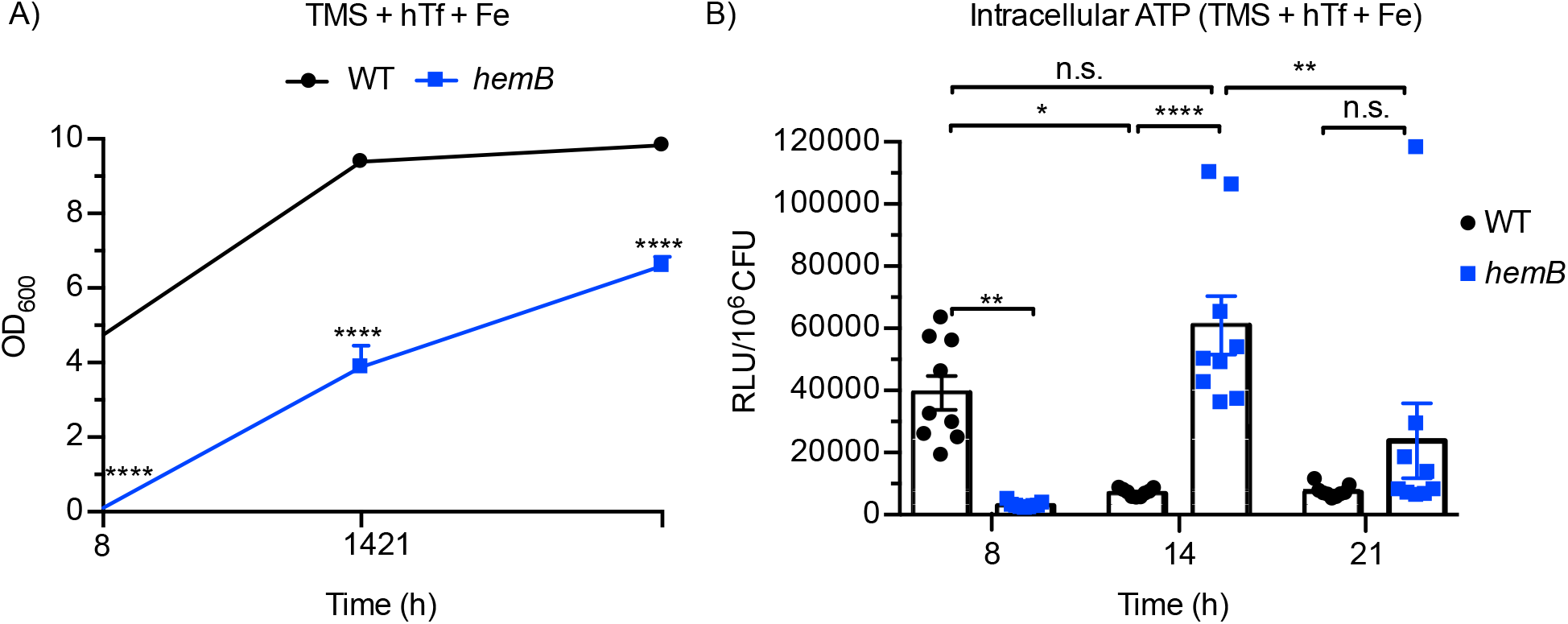
Intracellular ATP levels in a *S. aureus hemB* mutant in iron replete TMS. Growth **(A)** and intracellular ATP levels **(B)** of WT and *hemB S. aureus* grown in TMS supplemented with 0.4 µM hemin, 1.5 µM hTf, and 10 µM ferric ammonium sulfate (Fe). Data are plotted as mean ± SEM, nine biological replicates from three independent experiments. * *p* ≤ 0.05, ** *p* ≤ 0.01, **** *p* ≤ 0.0001, two-way ANOVA with Dunnett’s multiple comparisons **(A)**, one-way ANOVA with Tukey’s post-test **(B)**.

## References

1. Turner NA, Sharma-Kuinkel BK, Maskarinec SA, Eichenberger EM, Shah PP, Carugati M, et al. Methicillin-resistant *Staphylococcus aureus*: an overview of basic and clinical research. Nat Rev Microbiol. 2019;17: 203–218. doi:10.1038/s41579-018-0147-4

2. Tong SYC, Davis JS, Eichenberger E, Holland TL, Fowler VG. *Staphylococcus aureus* infections: Epidemiology, pathophysiology, clinical manifestations, and management. Clin Microbiol Rev. 2015;28: 603–661. doi:10.1128/CMR.00134-14

3. Boucher H, Miller LG, Razonable RR. Serious infections caused by methicillin-resistant *Staphylococcus aureus*. Clin Infect Dis. 2010;51 Suppl 2: S183–197. doi:10.1086/653519

4. Kochanek KD, Xu J, Murphy SL, Miniño AM, Kung H-C. Deaths: final data for 2009. Natl Vital Stat Reports. 2012;60: 1–116. Available: http://www.ncbi.nlm.nih.gov/pubmed/24974587

5. van Hal SJ, Jensen SO, Vaska VL, Espedido BA, Paterson DL, Gosbell IB. Predictors of mortality in *Staphylococcus aureus* bacteremia. Clin Microbiol Rev. 2012;25: 362–386. doi:10.1128/CMR.05022-11

6. Klevens RM, Morrison MA, Nadle J, Petit S, Gershman K, Ray S, et al. Invasive methicillin- resistant *Staphylococcus aureus* infections in the United States. J Am Med Assoc. 2007;298: 1763–1771. doi:10.1001/jama.298.15.1763

7. Prince A, Fok Lung TW. Consequences of Metabolic Interactions during *Staphylococcus aureus* Infection. Toxins (Basel). 2020;12. doi:10.3390/toxins12090581

8. Richardson AR. Virulence and Metabolism. Microbiol Spectr. 2019;7: 3–11. doi:10.1128/microbiolspec.GPP3-0011-2018

9. Dastgheyb SS, Otto M. Staphylococcal adaptation to diverse physiologic niches: An overview of transcriptomic and phenotypic changes in different biological environments. Future Microbiol. 2015;10: 1981–1995. doi:10.2217/fmb.15.116

10. Cassat JE, Skaar EP. Metal ion acquisition in *Staphylococcus aureus*: overcoming nutritional immunity. Semin Immunopathol. 2012;34: 215–235. doi:10.1007/s00281-011-0294-4

11. Somerville GA, Proctor RA. At the crossroads of bacterial metabolism and virulence factor synthesis in Staphylococci. Microbiol Mol Biol Rev. 2009;73: 233–248. doi:10.1128/MMBR.00005-09

12. Flannagan RS, Heit B, Heinrichs DE. Antimicrobial mechanisms of macrophages and the immune evasion strategies of *Staphylococcus aureus*. Pathogens. 2015;4: 826–868. doi:10.3390/pathogens4040826

13. Thammavongsa V, Kim HK, Missiakas D, Schneewind O. Staphylococcal manipulation of host immune responses. Nat Rev Microbiol. 2015;13: 529–543. doi:10.1038/nrmicro3521

14. Greenlee-Wacker M, DeLeo FR, Nauseef WM. How methicillin-resistant *Staphylococcus aureus* evade neutrophil killing. Curr Opin Hematol. 2015;22: 30–35. doi:10.1097/MOH.0000000000000096

15. Okumura CYM, Nizet V. Subterfuge and sabotage: Evasion of host innate defenses by invasive gram-positive bacterial pathogens. Annu Rev Microbiol. 2014;68: 439–458. doi:10.1146/annurev-micro-092412-155711

16. Krishna S, Miller LS. Host-pathogen interactions between the skin and *Staphylococcus aureus*. Curr Opin Microbiol. 2012;15: 28–35. doi:10.1016/j.mib.2011.11.003

17. Griffiths E. Iron in biological systems. In: Bullen JJ, Griffiths E, editors. New York: John Wiley & Sons, Ltd.; 1999. pp. 1–26.

18. Cassat JE, Skaar EP. Iron in infection and immunity. Cell Host Microbe. 2013;13: 509–519. doi:10.1016/j.chom.2013.04.010

19. Ganz T. Iron in innate immunity: starve the invaders. Current Opinion in Immunology Feb, 2009 pp. 63–67. doi:10.1016/j.coi.2009.01.011

20. Ganz T. Hepcidin and its role in regulating systemic iron metabolism. Hematol Am Soc Hematol Educ Progr. 2006; 29–35, 507. doi:10.1182/asheducation-2006.1.29

21. Weinberg ED. Nutritional immunity: Host’s attempt to withhold iron from microbial invaders. JAMA. 1975;231: 39–41. doi:10.1001/jama.1975.03240130021018

22. Sheldon JR, Laakso HA, Heinrichs DE. Iron Acquisition Strategies of Bacterial Pathogens. Microbiol Spectr. 2016;4: 43–85. doi:10.1128/microbiolspec.VMBF-0010-2015

23. Pinochet-Barros A, Helmann JD. Redox sensing by Fe^2+^ in bacterial Fur family metalloregulators. Antioxid Redox Signal. 2018;29: 1858–1871. doi:10.1089/ars.2017.7359

24. Sheldon JR, Heinrichs DE. Recent developments in understanding the iron acquisition strategies of gram positive pathogens. Whitfield C, editor. FEMS Microbiol Rev. 2015;39: 592–630. doi:10.1093/femsre/fuv009

25. Conroy BS, Grigg JC, Kolesnikov M, Morales LD, Murphy MEP. *Staphylococcus aureus* heme and siderophore-iron acquisition pathways. BioMetals. 2019;32: 409–424. doi:10.1007/s10534-019-00188-2

26. Tiedemann MT, Heinrichs DE, Stillman MJ. Multiprotein heme shuttle pathway in *Staphylococcus aureus*: iron-regulated surface determinant cog-wheel kinetics. J Am Chem Soc. 2012;134: 16578–85. doi:10.1021/ja305115y

27. Grigg JC, Ukpabi G, Gaudin CFM, Murphy MEP. Structural biology of heme binding in the *Staphylococcus aureus* Isd system. J Inorg Biochem. 2010;104: 341–348. doi:10.1016/j.jinorgbio.2009.09.012

28. Muryoi N, Tiedemann MT, Pluym M, Cheung J, Heinrichs DE, Stillman MJ. Demonstration of the iron-regulated surface determinant (Isd) heme transfer pathway in *Staphylococcus aureus*. J Biol Chem. 2008;283: 28125–36. doi:10.1074/jbc.M802171200

29. Maresso AW, Schneewind O. Iron acquisition and transport in *Staphylococcus aureus*. Biometals. 2006;19: 193–203. Available: http://www.ncbi.nlm.nih.gov/entrez/query.fcgi?cmd=Retrieve&db=PubMed&dopt=Citation&list_uids=16718604

30. Mazmanian SK, Skaar EP, Gaspar AH, Humayun M, Gornicki P, Jelenska J, et al. Passage of heme-iron across the envelope of *Staphylococcus aureus*. Science. 2003;299: 906–909. doi:10.1126/science.1081147

31. Pishchany G, Sheldon JR, Dickson CF, Alam MT, Read TD, Gell DA, et al. IsdB-dependent hemoglobin binding is required for acquisition of heme by *Staphylococcus aureus*. J Infect Dis. 2014;209: 1764–72. doi:10.1093/infdis/jit817

32. Hammer ND, Skaar EP. Molecular mechanisms of *Staphylococcus aureus* iron acquisition. Annu Rev Microbiol. 2011;65: 129–147. doi:10.1146/annurev-micro-090110-102851

33. Proctor RA, von Eiff C, Kahl BC, Becker K, McNamara P, Herrmann M, et al. Small colony variants: a pathogenic form of bacteria that facilitates persistent and recurrent infections. Nat Rev Microbiol. 2006;4: 295–305. doi:10.1038/nrmicro1384

34. Proctor RA, Kriegeskorte A, Kahl BC, Becker K, Löffler B, Peters G. Staphylococcus aureus Small Colony Variants (SCVs): a road map for the metabolic pathways involved in persistent infections. Front Cell Infect Microbiol. 2014;4: 99. doi:10.3389/fcimb.2014.00099

35. Kriegeskorte A, Grubmüller S, Huber C, Kahl BC, von Eiff C, Proctor RA, et al. *Staphylococcus aureus* small colony variants show common metabolic features in central metabolism irrespective of the underlying auxotrophism. Front Cell Infect Microbiol. 2014;4: 141. doi:10.3389/fcimb.2014.00141

36. Tuchscherr L, Löffler B, Proctor RA. Persistence of *Staphylococcus aureus*: Multiple metabolic pathways impact the expression of virulence factors in Small-Colony Variants (SCVs). Front Microbiol. 2020;11: 1028. doi:10.3389/fmicb.2020.01028

37. Kahl BC, Becker K, Löffler B. Clinical significance and pathogenesis of Staphylococcal small colony variants in persistent infections. Clin Microbiol Rev. 2016;29: 401–427. doi:10.1128/CMR.00069-15

38. Junge S, Görlich D, den Reijer M, Wiedemann B, Tümmler B, Ellemunter H, et al. Factors associated with worse lung function in cystic fibrosis patients with persistent *Staphylococcus aureus*. PLoS One. 2016;11: e0166220. doi:10.1371/journal.pone.0166220

39. Kahl BC, Duebbers A, Lubritz G, Haeberle J, Koch HG, Ritzerfeld B, et al. Population dynamics of persistent *Staphylococcus aureus* isolated from the airways of cystic fibrosis patients during a 6-year prospective study. J Clin Microbiol. 2003;41: 4424–4427. doi:10.1128/jcm.41.9.4424-4427.2003

40. Hammer ND, Reniere ML, Cassat JE, Zhang Y, Hirsch AO, Hood MI, et al. Two heme- dependent terminal oxidases power *Staphylococcus aureus* organ-specific colonization of the vertebrate host. MBio. 2013;4: 1–9. doi:10.1128/mBio.00241-13

41. Proctor RA, van Langevelde P, Kristjansson M, Maslow JN, Arbeit RD. Persistent and relapsing infections associated with small-colony variants of *Staphylococcus aureus*. Clin Infect Dis. 1995;20: 95–102. doi:10.1093/clinids/20.1.95

42. Von Eiff C, Heilmann C, Proctor RA, Woltz C, Peters G, Götz F. A site-directed *Staphylococcus aureus hemB* mutant is a small-colony variant which persists intracellularly. J Bacteriol. 1997;179: 4706–4712. doi:10.1128/jb.179.15.4706-4712.1997

43. Torres VJ, Stauff DL, Pishchany G, Bezbradica JS, Gordy LE, Iturregui J, et al. A *Staphylococcus aureus* regulatory system that responds to host heme and modulates virulence. Cell Host Microbe. 2007;1: 109–119. doi:10.1016/j.chom.2007.03.001

44. Stauff DL, Torres VJ, Skaar EP. Signaling and DNA-binding activities of the *Staphylococcus aureus* HssR-HssS two-component system required for heme sensing. J Biol Chem. 2007;282: 26111–26121. doi:10.1074/jbc.M703797200

45. Stauff DL, Skaar EP. The heme sensor system of *Staphylococcus aureus*. Contrib Microbiol. 2009;16: 120–35. doi:10.1159/000219376

46. Kohler C, von Eiff C, Liebeke M, McNamara PJ, Lalk M, Proctor RA, et al. A defect in menadione biosynthesis induces global changes in gene expression in *Staphylococcus aureus*. J Bacteriol. 2008;190: 6351–6364. doi:10.1128/JB.00505-08

47. Seggewiss J, Becker K, Kotte O, Eisenacher M, Yazdi MRK, Fischer A, et al. Reporter metabolite analysis of transcriptional profiles of a *Staphylococcus aureus* strain with normal phenotype and its isogenic *hemB* mutant displaying the small-colony-variant phenotype. J Bacteriol. 2006;188: 7765–7777. doi:10.1128/JB.00774-06

48. Moisan H, Brouillette E, Jacob CL, Langlois-Bégin P, Michaud S, Malouin F. Transcription of virulence factors in *Staphylococcus aureus* small-colony variants isolated from cystic fibrosis patients is influenced by SigB. J Bacteriol. 2006;188: 64–76. doi:10.1128/JB.188.1.64-76.2006

49. Kahl BC, Belling G, Becker P, Chatterjee I, Wardecki K, Hilgert K, et al. Thymidine-dependent *Staphylococcus aureus* small-colony variants are associated with extensive alterations in regulator and virulence gene expression profiles. Infect Immun. 2005;73: 4119–4126. doi:10.1128/IAI.73.7.4119-4126.2005

50. Poudel S, Tsunemoto H, Seif Y, Sastry A V., Szubin R, Xu S, et al. Revealing 29 sets of independently modulated genes in *Staphylococcus aureus*, their regulators, and role in key physiological response. Proc Natl Acad Sci U S A. 2020;117: 17228–17239. doi:10.1073/pnas.2008413117

51. Kriegeskorte A, Block D, Drescher M, Windmüller N, Mellmann A, Baum C, et al. Inactivation of *thyA* in *Staphylococcus aureus* attenuates virulence and has a strong impact on metabolism and virulence gene expression. MBio. 2014;5: e01447–14. doi:10.1128/mBio.01447-14

52. Cooper JD, Hannauer M, Marolda CL, Briere L-AK, Heinrichs DE. Identification of a positively charged platform in *Staphylococcus aureus* HtsA that is essential for ferric staphyloferrin A transport. Biochemistry. 2014;53: 5060–9. doi:10.1021/bi500729h

53. Grigg JC, Cooper JD, Cheung J, Heinrichs DE, Murphy MEP. The *Staphylococcus aureus* siderophore receptor HtsA undergoes localized conformational changes to enclose staphyloferrin A in an arginine-rich binding pocket. J Biol Chem. 2010;285: 11162–71. doi:10.1074/jbc.M109.097865

54. Grigg JC, Cheung J, Heinrichs DE, Murphy MEP. Specificity of staphyloferrin B recognition by the SirA receptor from *Staphylococcus aureus*. J Biol Chem. 2010;285: 34579–34588. doi:10.1074/jbc.M110.172924

55. Beasley FC, Vinés ED, Grigg JC, Zheng Q, Liu S, Lajoie GA, et al. Characterization of staphyloferrin A biosynthetic and transport mutants in *Staphylococcus aureus*. Mol Microbiol. 2009;72: 947–963. doi:10.1111/j.1365-2958.2009.06698.x

56. Beasley FC, Marolda CL, Cheung J, Buac S, Heinrichs DE. *Staphylococcus aureus* transporters Hts, Sir, and Sst capture iron liberated from human transferrin by Staphyloferrin A, Staphyloferrin B, and catecholamine stress hormones, respectively, and contribute to virulence. Infect Immun. 2011;79: 2345–55. doi:10.1128/IAI.00117-11

57. Sebulsky MT, Hohnstein D, Hunter MD, Heinrichs DE. Identification and characterization of a membrane permease involved in iron-hydroxamate transport in *Staphylococcus aureus*. J Bacteriol. 2000;182: 4394–400. Available: http://www.ncbi.nlm.nih.gov/htbin-post/Entrez/query?db=m&form=6&dopt=r&uid=0010913070 http://jb.asm.org/cgi/content/full/182/16/4394 http://jb.asm.org/cgi/content/abstract/182/16/4394

58. Podkowa KJ, Briere L-AK, Heinrichs DE, Shilton BH. Crystal and solution structure analysis of FhuD2 from *Staphylococcus aureus* in multiple unliganded conformations and bound to ferrioxamine-B. Biochemistry. 2014;53: 2017–31. doi:10.1021/bi401349d

59. Arifin AJ, Hannauer M, Welch I, Heinrichs DE. Deferoxamine mesylate enhances virulence of community-associated methicillin resistant *Staphylococcus aureus*. Microbes Infect. 2014;16: 967–72. doi:10.1016/j.micinf.2014.09.003

60. Garcia LG, Lemaire S, Kahl BC, Becker K, Proctor RA, Denis O, et al. Antibiotic activity against small-colony variants of *Staphylococcus aureus*: review of in vitro, animal and clinical data. J Antimicrob Chemother. 2013;68: 1455–64. doi:10.1093/jac/dkt072

61. von Eiff C. *Staphylococcus aureus* small colony variants: a challenge to microbiologists and clinicians. Int J Antimicrob Agents. 2008;31: 507–10. doi:10.1016/j.ijantimicag.2007.10.026

62. Guérillot R, Kostoulias X, Donovan L, Li L, Carter GP, Hachani A, et al. Unstable chromosome rearrangements in *Staphylococcus aureus* cause phenotype switching associated with persistent infections. Proc Natl Acad Sci U S A. 2019;116: 20135–20140. doi:10.1073/pnas.1904861116

63. Becker K, Al Laham N, Fegeler W, Proctor RA, Peters G, Von Eiff C. Fourier-transform infrared spectroscopic analysis is a powerful tool for studying the dynamic changes in *Staphylococcus aureus* small-colony variants. J Clin Microbiol. 2006;44: 3274–3278. doi:10.1128/JCM.00847-06

64. Garcia LG, Lemaire S, Kahl BC, Becker K, Proctor RA, Denis O, et al. Pharmacodynamic evaluation of the activity of antibiotics against hemin- and menadione-dependent small-colony variants of *Staphylococcus aureus* in models of extracellular (broth) and intracellular (THP-1 monocytes) infections. Antimicrob Agents Chemother. 2012;56: 3700– 3711. doi:10.1128/AAC.00285-12

65. Tuchscherr L, Heitmann V, Hussain M, Viemann D, Roth J, von Eiff C, et al. *Staphylococcus aureus* small-colony variants are adapted phenotypes for intracellular persistence. J Infect Dis. 2010;202: 1031–1040. doi:10.1086/656047

66. Biswas L, Biswas R, Schlag M, Bertram R, Götz F. Small-colony variant selection as a survival strategy for *Staphylococcus aureus* in the presence of *Pseudomonas aeruginosa*. Appl Environ Microbiol. 2009;75: 6910–6912. doi:10.1128/AEM.01211-09

67. von Eiff C, McNamara P, Becker K, Bates D, Lei X-H, Ziman M, et al. Phenotype microarray profiling of *Staphylococcus aureus menD* and *hemB* mutants with the small-colony-variant phenotype. J Bacteriol. 2006;188: 687–693. doi:10.1128/JB.188.2.687-693.2006

68. Vaudaux P, Francois P, Bisognano C, Kelley WL, Lew DP, Schrenzel J, et al. Increased expression of clumping factor and fibronectin-binding proteins by *hemB* mutants of *Staphylococcus aureus* expressing small colony variant phenotypes. Infect Immun. 2002;70: 5428–5437. doi:10.1128/iai.70.10.5428-5437.2002

69. Choby JE, Skaar EP. Heme Synthesis and Acquisition in Bacterial Pathogens. J Mol Biol. 2016;428: 3408–28. doi:10.1016/j.jmb.2016.03.018

70. Kohler C, von Eiff C, Peters G, Proctor RA, Hecker M, Engelmann S. Physiological characterization of a heme-deficient mutant of *Staphylococcus aureus* by a proteomic approach. J Bacteriol. 2003;185: 6928–6937. doi:10.1128/JB.185.23.6928-6937.2003

71. Lamontagne Boulet M, Isabelle C, Guay I, Brouillette E, Langlois J-P, Jacques P-É, et al. Tomatidine is a lead antibiotic molecule that targets *Staphylococcus aureus* ATP synthase subunit C. Antimicrob Agents Chemother. 2018;62. doi:10.1128/AAC.02197-17

72. Tavares AFN, Nobre LS, Melo AMP, Saraiva LM. A novel nitroreductase of *Staphylococcus aureus* with S-nitrosoglutathione reductase activity. J Bacteriol. 2009;191: 3403–3406. doi:10.1128/JB.00022-09

73. Kobylarz MJ, Heieis GA, Loutet SA, Murphy MEP. Iron uptake oxidoreductase (IruO) uses a flavin adenine dinucleotide semiquinone intermediate for iron-siderophore reduction. ACS Chem Biol. 2017;12: 1778–1786. doi:10.1021/acschembio.7b00203

74. Loutet SA, Kobylarz MJ, Chau CHT, Murphy MEP. IruO is a reductase for heme degradation by IsdI and IsdG proteins in *Staphylococcus aureus*. J Biol Chem. 2013;288: 25749–25759. doi:10.1074/jbc.M113.470518

75. Hannauer M, Arifin AJ, Heinrichs DE. Involvement of reductases IruO and NtrA in iron acquisition by *Staphylococcus aureus*. Mol Microbiol. 2015;96: 1192–210. doi:10.1111/mmi.13000

76. Skaar EP, Humayun M, Bae T, DeBord KL, Schneewind O. Iron-source preference of *Staphylococcus aureus* infections. Science. 2004;305: 1626–1628. Available: http://www.ncbi.nlm.nih.gov/pubmed/15361626

77. Andrews SC, Shipley D, Keen JN, Findlay JBC, Harrison PM, Guest JR. The haemoglobin-like protein (HMP) of *Escherichia coli* has ferrisiderophore reductase activity and its C-terminal domain shares homology with ferredoxin NADP+ reductases. FEBS Lett. 1992;302: 247– 52. doi:10.1016/0014-5793(92)80452-m

78. Laakso HA, Marolda CL, Pinter TB, Stillman MJ, Heinrichs DE. A heme-responsive regulator controls synthesis of staphyloferrin B in *Staphylococcus aureus*. J Biol Chem. 2016;291: 29– 40. doi:10.1074/jbc.M115.696625

79. Sheldon JR, Marolda CL, Heinrichs DE. TCA cycle activity in *Staphylococcus aureus* is essential for iron-regulated synthesis of staphyloferrin A, but not staphyloferrin B: the benefit of a second citrate synthase. Mol Microbiol. 2014;92: 824–39. doi:10.1111/mmi.12593

80. Perry WJ, Spraggins JM, Sheldon JR, Grunenwald CM, Heinrichs DE, Cassat JE, et al. *Staphylococcus aureus* exhibits heterogeneous siderophore production within the vertebrate host. Proc Natl Acad Sci U S A. 2019; 2–4. doi:10.1073/pnas.1913991116

81. Robins-Browne RM, Prpic JK. Desferrioxamine and systemic yersiniosis. Lancet (London, England). 1983;2: 1372. doi:10.1016/s0140-6736(83)91136-4

82. Skaar EP, Humayun M, Bae T, DeBord KL, Schneewind O. Iron-source preference of *Staphylococcus aureus* infections. Science. 2004;305: 1626–1628. doi:10.1126/science.1099930

83. Hammer ND, Cassat JE, Noto MJ, Lojek LJ, Chadha AD, Schmitz JE, et al. Inter- and intraspecies metabolite exchange promotes virulence of antibiotic-resistant *Staphylococcus aureus*. Cell Host Microbe. 2014;16: 531–537. doi:10.1016/j.chom.2014.09.002

84. Jonsson I-M, von Eiff C, Proctor RA, Peters G, Rydén C, Tarkowski A. Virulence of a *hem*B mutant displaying the phenotype of a *Staphylococcus aureus* small colony variant in a murine model of septic arthritis. Microb Pathog. 2003;34: 73–79. doi:10.1016/s0882-4010(02)00208-5

85. Brouillette E, Martinez A, Boyll BJ, Allen NE, Malouin F. Persistence of a *Staphylococcus aureus* small-colony variant under antibiotic pressure in vivo. FEMS Immunol Med Microbiol. 2004;41: 35–41. doi:10.1016/j.femsim.2003.12.007

86. Bates DM, von Eiff C, McNamara PJ, Peters G, Yeaman MR, Bayer AS, et al. *Staphylococcus aureus menD* and *hemB* mutants are as infective as the parent strains, but the menadione biosynthetic mutant persists within the kidney. J Infect Dis. 2003;187: 1654–1661. doi:10.1086/374642

87. Seilie ES, Bubeck Wardenburg J. *Staphylococcus aureus* pore-forming toxins: The interface of pathogen and host complexity. Semin Cell Dev Biol. 2017;72: 101–116. doi:10.1016/j.semcdb.2017.04.003

88. Sebulsky MT, Speziali CD, Shilton BH, Edgell DR, Heinrichs DE. FhuD1, a ferric hydroxamate- binding lipoprotein in *Staphylococcus aureus*: a case of gene duplication and lateral transfer. J Biol Chem. 2004;279: 53152–9. doi:10.1074/jbc.M409793200

89. Dale SE, Sebulsky MT, Heinrichs DE. Involvement of SirABC in iron-siderophore import in *Staphylococcus aureus*. J Bacteriol. 2004;186: 8356–62. doi:10.1128/JB.186.24.8356- 8362.2004

90. Bateman BT, Donegan NP, Jarry TM, Palma M, Cheung a L. Evaluation of a tetracycline-inducible promoter in *Staphylococcus aureus* in vitro and in vivo and its application in demonstrating the role of *sigB* in microcolony formation. Infect Immun. 2001;69: 7851–7. doi:10.1128/IAI.69.12.7851-7857.2001

91. Bae T, Schneewind O. Allelic replacement in *Staphylococcus aureus* with inducible counter- selection. Plasmid. 2006;55: 58–63. doi:10.1016/j.plasmid.2005.05.005

92. Love MI, Huber W, Anders S. Moderated estimation of fold change and dispersion for RNA-seq data with DESeq2. Genome Biol. 2014;15: 550. doi:10.1186/s13059-014-0550-8

